# Nucleus raphe magnus serotonin neurons bidirectionally control spinal nociceptive transmission in mice

**DOI:** 10.1101/2025.04.09.647804

**Authors:** Grivet Zoé, Verboven Aude, Aby Franck, Bouali-Benazzouz Rabia, Dhellemmes Thibault, Perrot Emma, Beyeler Anna, Lora K Heisler, Balia Maddalena, Battefeld Arne, Martin Hugo, Retailleau Aude, Maingret François, Fioramonti Xavier, Landry Marc, De Koninck Yves, Benazzouz Abdelhamid, Fossat Pascal

## Abstract

**Background:** Noxious stimuli are conveyed to and integrated in the dorsal horn of the spinal cord (DHSC) before being transmitted to supraspinal centers, where pain perception is generated. Descending pathways from the brainstem dynamically modulate this integration, either facilitating or inhibiting nociceptive information based on physiological, emotional, genetic and environmental factors. Serotonergic neurons in the nucleus raphe magnus (NRM), activating different spinal 5-HT receptors, exert bidirectional control, both facilitatory and inhibitory, but the underlying mechanisms of this control remain unclear.

**Methods:** We investigated in adult mice, the NRM serotonergic modulation of nociception using imaging, behavioral, pharmacological, electrophysiological, chemogenetic and optogenetic approaches.

**Results:** We have demonstrated that the action of serotonergic neurons in the NRM on spinal nociceptive transmission depends on their activation pattern, which targets different spinal 5-HT receptors likely associated with different spinal microcircuits. Serotonergic neurons of the NRM exert a tonic analgesic effect mediated by 5-HT_2c_ receptor. Low increase in 5-HT activity leads to an increased analgesia through spinal inhibitory interneurons expressing 5-HT_2c_ and 5-HT_2A_ receptors. Finally, prolonged stimulation of serotonergic neurons led to hyperalgesia mediated by 5-HT_3_ receptor. Comparison of 5-HT receptors in spinal tissue from mice and human shows that 5-HT_2c_ receptor has a very high expression level, comparable between both species.

**Conclusions:** These results propose a model of bidirectional action of serotonin neurons on nociceptive transmission depending on their level of activity and show that 5-HT_2c_ receptor is the main mediator of serotonin-induced analgesia.

## INTRODUCTION

Nociceptive signals convey harmful information to several regions of the brain that generate the conscious perception of pain^1^. Before this multidimensional sensation emerges, these nociceptive signals are transmitted from peripheral tissues to the dorsal horn of the spinal cord (DHSC), where they are integrated and modulated^2^. This integration is tightly regulated by descending pathways originating from the brainstem, which can either amplify or suppress nociceptive signaling depending on physiological, emotional, and environmental factors ^2–5^. The interplay between these opposing influences ensures that nociception is precisely tuned, allowing pain to serve as a timely non-pathological alarm signal.

Among descending modulatory pathways, serotonergic (5-hydroxytryptamine; 5-HT) neurons of the nucleus raphe magnus (NRM) were among the first to be identified as key players in pain modulation in the 1970s ^6–8^. The translational relevance of this discovery is illustrated by the effectiveness of 5-HT and noradrenaline reuptake inhibitors currently prescribed to treat neuropathic pain ^9^. Early models proposed an inhibitory role for 5-HT pathways mediated by activation of spinal opioid microcircuits^6^. However, the discovery of 14 different subtypes of 5-HT receptors expressed in the spinal cord, revealed a more complex picture, with 5-HT exerting both inhibitory and excitatory effects depending on the concentration of 5-HT and/or the 5-HT receptor activated (For review see^10–12^). Spinal application of 5-HT induces bidirectional, either excitatory or inhibitory action on DHSC neurons and nociceptive behavior^13–16^. The dose of 5-HT used can modify the behavioral outcomes suggesting that the level of activity of 5-HT neurons can influence the effect on nociceptive transmission^17^. 5-HT descending control is likely due to a dose-dependent action of 5-HT on one or more of the fourteen 5-HT receptors with analgesia mainly mediated via 5-HT_1_ receptor family (e.g. 5-HT_1A_) and proalgesia associated with the ionotropic 5-HT_3_ receptor^4,11,18–22^. Most of these studies used either 5-HT or related modulators administered spinally, so the link between brainstem 5-HT neuron activity and spinal nociceptive transmission is indirect.

Recent technological advances including optogenetics and chemogenetics associated with the development of transgenic mice have enabled precise targeting of 5-HT neuronal populations of the brainstem projecting to the DHSC. Once again, the dual action of 5-HT have been observed with either a pronociceptive or an analgesic effects^23–25^. A critical challenge in explaining these seemingly contradictory findings, lies in the variability of experimental paradigms, particularly differences in the level and duration of 5-HT neuron activation, which complicate direct comparisons between studies. Our hypothesis is that brainstem 5-HT neurons activity tune the direction of spinal nociceptive transmission. To test this hypothesis, we used optogenetic and chemogenetic approaches to specifically manipulate 5-HT neurons of NRM (5-HT^NRM^) and changed the parameters of stimulation to observe the consequences on spinal nociceptive transmission. We report that 5-HT^NRM^ exerts, on the one hand, a tonic inhibition of nociceptive transmission that is reinforced by low level of 5-HT^NRM^ activity and mediated through activation of spinal inhibitory interneurons by 5-HT2_A/C_ receptors. On the other hand, a high level of 5-HT^NRM^ activity leads to an increase in nociceptive transmission mediated by 5-HT_3_ receptors probably located on excitatory neurons in the DHSC.

## Methods

### Animals

Adult (20-25g) female and male wild type (C57BL/6J; Janvier labs, France) or *ePet-Cre* mice (B6.Cg-Tg (FeV-cre) 1Esd/J, Jackson Laboratory, USA) were used. Heterozygous *ePet-Cre* mice are transgenic animals that express the cre-recombinase under the control of the Pet-1 enhancer (*ePet*), a sequence allowing specific expression of PET-1 protein strictly restricted to 5-HT synthetizing neurons ^26^. *ePet-Cre* were crossbred with Ai9tdtomato line mice (Gt(Rosa)26Sortm6 (CAG-tdTomato)Hze, Jackson Laboratory) in order to obtain, *ePet-Cre*::Ai9tdtomato mice. Heterozygous *Htr2a-Cre::Ai14* (*5-HT_2A_-Cre::Ai14)* and homozygous *Htr2c-Cre::Ai14* (*5-HT_2C_-Cre::Ai14*), mouse lines expressing tdtomato under the control of the CRE recombinase expressed either in cells expressing 5-HT_2c_ or 5-HT_2A_ receptors were also used.

All mice had free access to food and water. They were housed in cages of 5 animals and maintained at constant temperature (21±2°C) and humidity levels (60%), with a 12/12 hour light/dark cycle. All cages were enriched with a red plastic tunnel, wooden sticks, cotton squares and bedding (litter). All procedures have been approved by the ethical committee of Bordeaux and the Ministry of research (CE50; Animal Facility PIV-EXPE, APAFIS#25605, APAFIS#32137 and APAFIS#26108; Animal Facilities of Neurocentre Magendie, APAFIS#21068). All efforts have been made to minimize animal suffering and to reduce the number of animals used.

### Surgery

Mice were anaesthetized under 4% isoflurane in an induction chamber then were placed into a stereotactic frame (RWD, Shengzen, China) on a heating pad, head-fixed with ear bars and maintained at 1.5-2% isoflurane. The eyes were protected with ocular gel and the area of interest shaved and then cleaned with betadine. Mice received a subcutaneous injection of buprenorphine (100µl, 0.1mg/kg) and local injection of lidocaine (100µl, 0.4mg/kg). Injections were made using a graduated glass capillary with an outer tip diameter of 40µm attached to a plastic tube and a 5ml syringe.

For optogenetic and chemogenetic, we proceed to stereotactic injections of viruses in the nucleus raphe magnus (NRM) as previously described^27,28^ **(see Table S1)**. We injected a total of 600nl (in 3 boluses of 200nl, at coordinates from bregma (mm) antero-posterior/medio-lateral/dorso-ventral -5.6/0/-5.6; -5.8/0/-5.6; -6.1/0/-5.7), of selected viruses. Special care was then provided for 2 weeks before the implantation of the optic fibers in the optogenetic animal group.

Optical fibers were prepared and cleaved to the appropriate length (0.5mm) then polished, and their ability to transmit light was recorded using a Thorlabs power meter. Optic fiber implantations were performed 2 weeks after injection of viruses. Optical cannula (KFP2301LZXX LC Ceramic Ferule NO flange OD 1.25mm - ID 230μm - conc < 20μm from AMS Technologies, associated with Ø200μm Core TECS-Clad Multimode Optical Fiber, 0.39NA from Thorlabs) were implanted above the surface of the DHSC to deliver light to 5-HT descending fibers expressing the selected opsins.

Spinal cord implantations are performed according to the protocol described in^25,29,30,31^. In anesthetized mice (see above “surgery”). A 1- to 2-cm incision was made slightly caudal to the peak of the dorsal hump to expose the lumbar spinal region. The T12-L2 vertebra of interest was identified, and a small incision was made between the tendons and the vertebral column on either side. The vertebra was then secured using spinal adaptor clamps, and all the tissue was removed from the surface of the bone. Using a microdrill, we removed the spinous process and the surface of the vertebra. Next, a small hole was drilled approximately 1mm from midline, centered on the rostro-caudal axis on the left side. We positioned the optical fiber above the hole and cemented the cannula in place using first cyanoacrylate glue then dental cement. Finally, we sutured and cleaned the skin surrounding the dental cement, securing the cannula implantation. We monitored the animal for at least 1 week before behavioral and electrophysiological experiments.

### Pharmacology

#### DREADD

Deschloroclozapine dihydrochloride (DCZ) (water soluble) (0.1mg/kg, HelloBio (HB9126)) was freshly weighed and dissolved in saline (NaCl 0.9%) and intraperitoneally injected.

#### 5-HT receptors antagonists (see Table S2)

Application of antagonists was performed by injecting 10 µL solution containing different 5-HT receptor antagonists (see table) at the level of the spinal cord via an intrathecal injection followed by 5µl saline (NaCl 0.9%). Intrathecal injections were performed with a 10µl Hamilton syringe equipped with a 30G needle. The needle was inserted in the subarachnoid space between vertebrae L5 and L6. Drugs were dissolved either in NaCl 0.9% solution or in DMSO 1%. Solubilisation in DMSO was performed at high concentration and the final concentration was obtained by diluting with water. Control experiments with 10µl of the different solvent (NaCl or DMSO) alone were also performed.

### Nociceptive responses

Behavioral experiments were carried out blinded to experimental conditions. For optogenetic studies, the experimenter was blind to the virus (ChR2 or GFP) injected. For pharmacology studies, the experimenter was blind to the substance (drug or vehicle) injected.

Experimental designs are provided in **SupFig 1**.

Paw withdrawal threshold (PWT). Animals were placed in a square plastic frame with a mesh grid on the floor. After 10min habituation, animals were tested for PWT using Von Frey filaments (Bioseb, France). Five successive tests were performed by applying Von Frey filaments on the plantar surface of the paw in mice remaining on their four paws. We assessed the paw withdrawal threshold (PWT) using the simplified up and down (SUDO) technique^25,29,32^. Briefly, starting from the 10th filament (2g), the response of the animal to the stimulation (withdrawal or not) will decide the value of the next filament. The lower filament is tested if the animal withdraws the paw and the upper one in the other case. Five successive filaments are applied. Each test is separated by around 20s. A series of SUDO was performed before, during and after optogenetic stimulation of 5-HT neurons of the NRM or spinal cord. PWT was evaluated using the last filament value +/- 0.5 depending on the value of the fourth filament. To determine the value in gram, the algorithm PWT = 10^(x*F+B)^ was used. F is the filament number obtained using SUDO and x and B were determined from a linear regression of the logarithm of the empirically measured filament bending force plotted against the filament number (the following equation was used: Log (bending force) = x * Filament number + B, with x = 0.182 and B=-1.47 when 7<F<14, and x=0.240 and B=-2 when 2<F<9).

Repetitive stimulations (Number of responses). To assess the analgesic effect, after determining the PWT, a supra-threshold filament was selected, and five repetitive filament applications were performed before, during, and after optogenetic stimulation of the 5-HT neurons. The number of paw withdrawals was plotted as a percentage of the total number of applications (i.e., five). To test hyperalgesia effects, the same procedure was applied but with the choice of a subthreshold filament.

Thermal latency (paw withdrawal latency). Animals were placed in a Plexiglas cage with a transparent glass floor. An infrared laser beam of calibrated value was applied on the plantar surface of the paw until the animal withdrew its paw (Ugo Basile, Gemonio, Italy). The value of latency was measured in seconds. Two trials separated by at least 30s to avoid sensitization were performed before, during, and after optogenetic stimulation of 5-HT neurons of the NRM. The average value of the two trials was then compared between conditions. Laser intensity was adapted to the pathophysiological condition: IR50 (infrared intensity, internal calibration of the device) was used for naïve animals that give a mean withdrawal latency around 5 to 10s. A cut off of 20s was applied to avoid paw damages.

### *In vivo* electrophysiology

*In vivo* electrophysiology was performed on a subset of mice during a period of one week following the end of behavioral tests as previously described^25,29^ (see **SupFig1**). Mice were anesthetized with urethane 20% and placed on a stereotactic frame (M2E, Asnières, France). A laminectomy was performed on lumbar vertebrae T12–L2 and segments L4–L5 of the spinal cord were exposed. Extracellular recordings of second order DHSC neurons were made with borosilicate glass capillaries (2 MΩ, filled with NaCl 684mM) (Harvard Apparatus, Cambridge, MA, USA). For single unit recordings, the signal was high pass filtered and the criterion for the selection of a neuron was the presence of a A-fiber response (0-80ms) followed by a c-fiber response (80 to 300ms) to electrical stimulations of the ipsilateral paw with bipolar subcutaneous stimulation electrodes (single shock, 0-5mA, 1ms duration, AMPI, iso-flex stimulator, Israel). First, we determined the minimal stimulation allowing a delayed response of the neuron, i.e., an action potential between 80 and 300ms. We then set the intensity at 3 times the threshold and stimulated the paws again every 30s. The response of the neurons before and during optogenetic stimulation was recorded. Finally, we assessed the wind-up, a form of central sensitization. Windup protocol consisted of recording the response of DHSC neurons to 15 stimulations at 1Hz and at 3 times the threshold. A windup coefficient was calculated as the sum of c-spikes following each stimulation subtracted by 15 times the number of spikes following the first stimulation.

### Patch clamp recordings

Patch-clamp recordings were performed on brain slices from *ePet cre::Ai9td tomato* mice or ePet cre expressing either ArchT following injection of an AAV-DIO-ArchT-EYFP as previously described^25,29^. Briefly, the virus was injected in 8-week-old mice, and the patch clamp was applied when they were 11 to 12 weeks old. The mice were perfused intracardially during euthanasia (exagon/lidocaine: 300/30mg/kg*, i.p.*) with ice-cold N-methyl-D-glucamine (NMDG) solution (containing in mM: 1.25 NaH_2_PO_4_, 2.5 KCl, 7 MgCl_2_, 20 HEPES, 0.5 CaCl_2_, 28 NaHCO_3_, 8 D-glucose, 5L(+)-ascorbate, 3 Na-pyruvate, 2 thiourea, 93 NMDG, and 93 HCl 37%; pH: 7.3–7.4; osmolarity: 305–310mOsM). Brains were quickly removed and 250μm slices containing the NRM were prepared with a VT1100S Leica vibratome in ice-cold oxygenated NMDG solution before recovery for 12-15 minutes at 34°C in oxygenated NMDG solution. Slices were then transferred at room temperature into aCSF solution (containing in mM: 124 NaCl, 2.5 KCl, 1.25 NaH_2_PO4, 2 MgCl_2_, 2.5 CaCl_2_, 12.5 D-glucose, 25 NaHCO_3_; pH: 7.3-7.4; osmolarity: 305-310mOsM) for at least 1 hour. Slices were placed in the recording chamber under a microscope (Nikon EF600) outfitted for fluorescence and IR-DIC video microscopy and perfused with oxygenated aCSF at 2-3ml/min. Viable 5-HT^NRM^ neurons were identified with fluorescence and a video camera (Nikon). Borosilicate pipette (4-6MΩ; 1.5mm OD, Sutter Instrument) were filled with an intracellular solution (containing in mM: 128 Kgluconate, 20 NaCl, 1 MgCl_2_, 1 EGTA, 0.3 CaCl_2_, 2 Na_2_-ATP, 0.3 Na-GTP, 0.2 cAMP, 10 HEPES; 280-290 mOsM, pH 7.3-7.4). Recordings were made using a Multiclamp 700B amplifier, digitized using the Digidata 1440A interface and acquired at 2kHz using pClamp 10.5 software (Axon Instruments, Molecular Devices, Sunnyvale, CA). Pipette and cell capacitances were fully compensated but junction potential was not corrected. 5-HT^NRM^ neurons were recorded in whole-cell current-clamp mode. Brain slices expressing ArchT were opto-stimulated at 561nm (200ms continuous pulse) at a laser intensity of 5mW.mm^2^ and action potential frequency was recorded before, during and after optogenetic stimulation.

### Optogenetic stimulation and c-Fos experiments

ePet-cre mice expressing ChR2 in the NRM (see above) were anesthetized with urethane 20% and placed on a stereotactic frame (M2E, Asnières, France). Three group of 4 mice were prepared as follow: ChR2, GFP and Naive (without viral injection). Surgical procedure was performed as previously described (see above). Briefly, a laminectomy was performed on lumbar vertebrae T12 to L2, and segments L4 to L5 of the spinal cord were exposed. An optic fiber was placed above the spinal cord to allow light illumination of 5-HT descending fibers. After 1h of stable anesthesia, an optogenetic stimulation at 5Hz/5ms for 2min was performed. One hour later, mice were quickly euthanized and perfused with 4% (w/v) paraformaldehyde (PFA) in 0.1M phosphate-buffered saline (PBS). Spinal cords were extracted. dissected out, post-fixed for 3h at 4°C in the same solution, then cryoprotected in a solution of 0.1M PBS and 25% sucrose overnight, and stored at −80°C. c-Fos immunostaining with rabbit anti-c-Fos antibody (1/500; 9F6, mAB Cell Signalling Technology 2250 S) was performed as below (see immunohistochemistry section).

### Immunohistochemistry

Mice, used in *in vivo* experiments, were kept deeply anaesthetized with urethane 20% and then transcardially perfused with 50ml of NaCl 0.9%, Heparin (25UI/5ml) and 50ml of 4% PFA. Both brain and spinal cord were dissected and fixed overnight in 4% PFA, and then cryopreserved in 12.5% phosphate buffer solution (PBS) sucrose. After being frozen in Tissue-Tek O.C.T, brain and spinal cord coronal sections were cut at 20µm with a cryostat (Leica CM3050S) and kept in PBS-Azide1X 0.2%. After 3 bath washes of PBS1X, sections were put free-floating in a solution of 0.1M PBS / bovine serum albumin (BSA) 1% / triton 0.3% to permeabilize and saturate the slices. Slices were then bathed overnight at 4°C in a solution composed of primary antibodies anti-TPH2 (rabbit, Biotechne NB300-109), anti-GFP (chicken, Aves GFP-1010), anti-Pax2 (goat, AF3364 bio-techne), anti-Tlx3 (guinea pig, kind gift by Dr Muller^33^), anti-c-Fos (rabbit, 1:500; 9F6, mAB Cell Signalling Technology 2250 S) / PBS1X / BSA 1%. After being washed in PBS1X, secondary antibodies were added in PBS1X and BSA 1%. Depending on the primary antibody species, we used Alexa Fluor 568–conjugated goat anti-guinea pig (1:500, Molecular Probes), Alexa Fluor 568–conjugated donkey anti-goat (1:500; Thermo Fisher Scientific), Alexa Fluor 568– or 647–conjugated goat anti-rabbit (1:500; Invitrogen), and associated with Alexa Fluor 488–conjugated goat anti-chicken (1:500; Thermo Fisher Scientific).

### RNAscope ISH procedure

RNAscope was performed over a three-day period following the manufacturer’s instructions and utilizing the RNAscope® Multiplex Fluorescent Detection Kit v2 (ACD, Newark, CA, USA; #323110). On the first day, frozen fresh brain tissues (n=3) were sectioned using a cryostat, and 16 μm fresh tissue sections were mounted directly onto slides. They were then incubated in cold 4% PFA (4°C) for 16 minutes. Subsequently, the sections were dehydrated in alcohol baths of 50%, 70%, and 100%, with each percentage bath lasting 5min. The sections were then transferred to another 100% ethanol bath and left overnight at -20°C. The following day, sections were allowed to air-dry at room temperature for 10min. A hydrophobic barrier was drawn along the sections using a hydrophobic pen (ImmEdge Hydrophobic Barrier Pen, #310018), and the sections were left at room temperature for 10 minutes to allow the barrier to dry. The sections were then treated with Protease Pretreat-4 (Cat#322340) for 20min at room temperature. Afterward, the sections were washed with Wash buffer (Cat#310091) diluted at 1/50 in 0.1M PBS. Then, the sections were incubated with a mixture of three probes for 2 hours at 40°C. The probes used target the following genes: SLC32A1(ref: 319191-C2), SLC17A6 (ref: 319171-C3), HTR2C (ref: 401001), HTR3A (ref: 411141) and HTR2A (ref: 401291) (biotechne, US). Triplex staining was performed with SLC32A1 as C2, SLC17A6 as C3 associated with one of the three above HTR probes as C1.

Two control sections were also prepared in parallel: one positive control and one negative control. The significance of RNAscope lies in its signal amplification, which occurs in several steps: the sections were incubated with AMP1 for 30min at 40°C, followed by AMP2 and AMP3. Then section area incubated with horseradish peroxidase HRP-C1 for 15min at 40°C, washed with Wash Buffer 1X then incubated with Tyramide Signal Amplification (TSA) fluorophore of the chosen color (647nm) for C1 for 30min at 40°C. HRP was then blocked with the incubation of the section in HRP blocker solution for 15min at 40°C. These steps were repeated with HRP-C2 and HRP-C3 with TSA of different color (c3=520nm; c2=570nm) each time.

Subsequently, the sections were incubated with DAPI (#320851) for 20min at room temperature. Finally, the sections were mounted with Fluoromount (ThermoFisher Scientific) and kept in the dark at room temperature overnight in the dark at 4°C to dry. The sections were stored at -20°C until confocal acquisitions were performed. Images were taken with automatized confocal microscope (Zeiss Cell Discoverer 7).

### 5-HT receptors spinal expression processing

The GEO datasets GSE190442 (8 humans single nucleus samples), GSE103840 (single cell mice), GSE 103892 (single nucleus mice) were processed with R (4.5.1) and Seurat. After importing the gene count matrices, normalisation of gene expression was performed across all 55289 nuclei/cells of each dataset. Neurons were re-grouped based on their “subtype annotation” metadata into dorsal excitatory and inhibitory neurons respectively. For GSE190442, 5-HT receptors expression was analyzed in 631 excitatory and 471 inhibitory dorsal neurons. All eight samples were included in the analysis and the averaged expression per gene calculated.

### Cell quantification

Except for the RNAscope analyses, images were acquired with a confocal microscope (Leica TCS SP5), equipped with oil immersion objectives varying from 20x to 63x magnification. Image stacks were obtained between 10µm and 20µm. Positive cells were quantified with the Cell Counting plugin of ImageJ software. For the RNAscope, quantification was performed with ImageJ using a script based on the 3D-OC plugin (https://www.protocols.io/view/bioloc3d-b3d-protocol-for-3d-automated-analysis-of-g6ambzac7.html). This script allowed the quantification of specific labelling and the colocalization between two markers, by separating pixels considered as part of an object or part of the background.

### Statistical analysis

Comparison between groups has been performed with Prism (Graphpad software, US). Depending on the experimental groups, the number of variables, and the individual distribution (normal or not), several statistical tests were performed. When several variables were tested (effect of stimulation in several experimental groups), a two-way ANOVA was performed. When several groups were tested for a variable on individuals not following a normal distribution, a Kruskal-Wallis test followed by a Dunn test was performed. When two related groups were tested, if the sample followed a normal distribution, a paired test was performed; otherwise, a Wilcoxon test was performed. p<0.05 was considered significant.

All detailed statistical tests are gathered in the supplementary table S3 for main figures and supplementary table S4 for supplemental figures.

## RESULTS

### 5-HTNRM neuron activation increases nociceptive sensitivity

In order to study the precise role of 5-HT^NRM^ neurons in nociceptive transmission *in vivo,* we manipulated these neurons using *ePet-Cre* transgenic mice combined with stereotaxic injection of cre-dependent viruses to express DREADD or opsins (see **SupFig2**). To do this, we injected the AAV8-hSyn-DIO-hM3DGq virus into the NRM in *ePet-Cre* mice, enabling specific expression of the hM3Dq (Gq) DREADD excitatory receptor (5-HT^NRM^::hM3Dq mice) or the control virus AAV5-CAG-Flex-eGFP-WPRE (5-HT^NRM^::eGFP mice) (**Figure 1A**). Von Frey test shows that selective activation of 5-HT^NRM^ neurons via DREADD ligand deschloroclozapine (DCZ; 0.1mg/kg, i.p.) produced a significant decrease in the paw withdrawal threshold (PWT) compared to saline treatment in 5-HT^NRM^::hM3Dq mice or to DCZ in 5-HT^NRM^::eGFP mice (**Figure 1B and SupFig3A-B**).

**Fig. 1.**
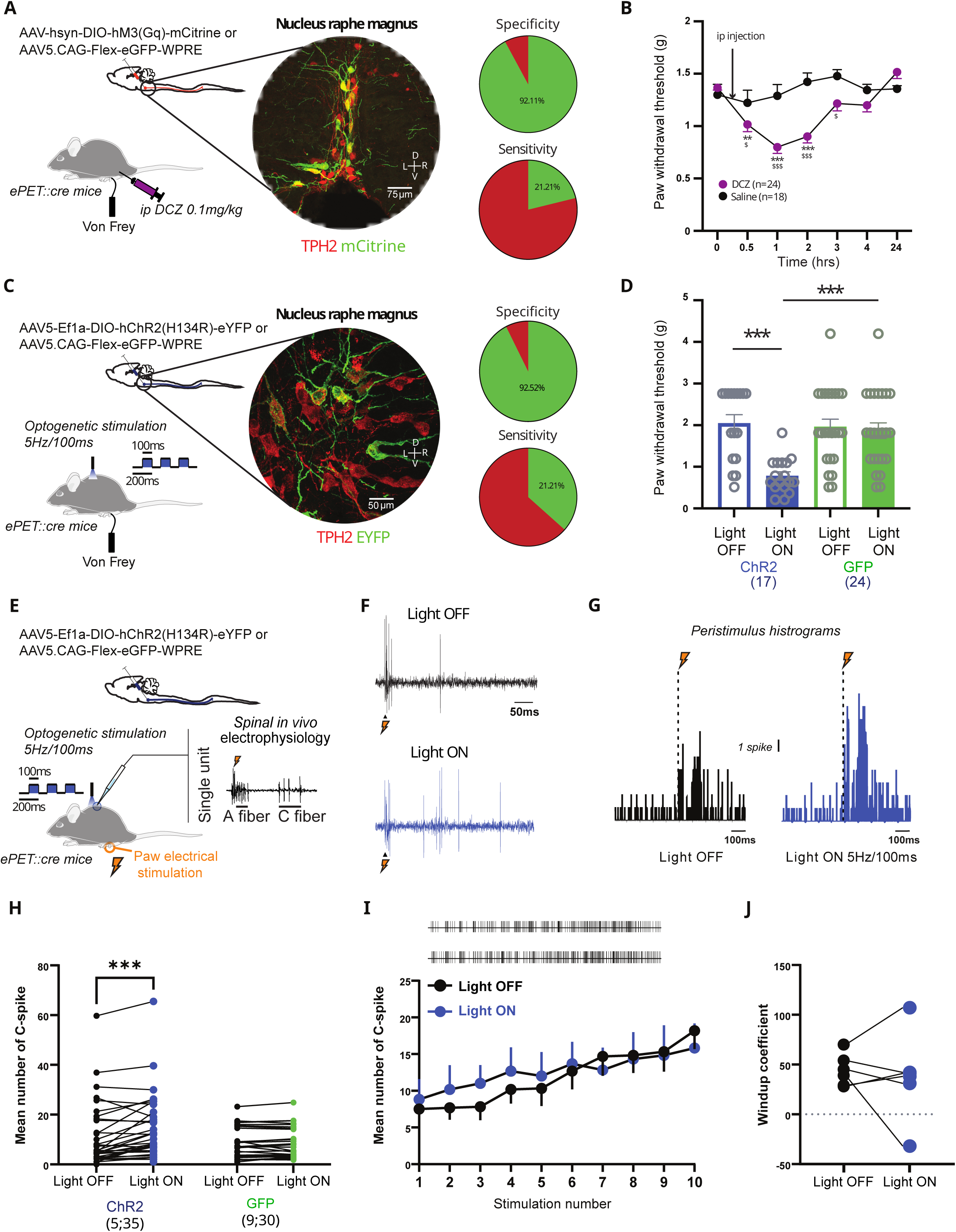
Long term activation of 5-HT neurons is proalgesic. (A) ePet-cre mice are injected with a cre-dependent AAV to express the excitatory DREADD hDM3. Activation of 5-HT neurons is done with ip injection of DCZ and PWT is assessed with Von frey filaments. Expression of mCitrine in the NRM colocalized with TPH2 with a high specificity. (B) ip injection of DCZ but not saline induced a mechanical allodynia for 2 hours (Two-way ANOVA, p<0.01, *vs 0 and $ DCZ vs saline). (C) ePet-cre mice are injected with a cre-dependent AAV to express the excitatory opsin ChR2. Activation of ChR2 is done through an optic fiber placed above the lumbar spinal cord at 5Hz/100ms. Expression of the eYFP colocalized with TPH2 showing high specificity. (D) Optogenetic activation of 5-HT fibers induce a decrease in PWT in the ChR2 group but not in GFP group alone (two-way ANOVA, p<0.0001). (E) single unit extracellular recordings of DHSC neurons in e-Pet mice injected with a cre-dependent AAV to express ChR2 in 5-HT neurons. (F) example of DHSC neuron response to supraliminar electric stimulation of the peripheral receptive field before (upper panel) and during optogenetic stimulation (lower panel). (G) Left; peristimulus histogram of DHSC neurons in response to suprathreshold electric shock on the receptive field in the hind paw. Right; peristimulus histogram of the same DHSC neurons in response to peripheral electric stimulation but during optogenetic stimulation of the 5-HT descending fibers. (H) On average, optogenetic stimulation of 5-HT neurons significantly increased the number of c-fiber mediated action potentials in ChR2 but not in GFP group (Two-way ANOVA, p<0.001).In brackets (number of mice; number of neurons). (I) Windup of DHSC neurons is not modified by optogenetic stimulation of 5-HT fibers (Two-way ANOVA, p>0.05). (J) Windup coefficient (Paired t test, p>0.05). Data are presented in mean ± SEM.

DREADD approach does not address the precise circuit involved, therefore, we next sought to clarify the neuronal circuit by focusing on 5-HT^NRM^ → DHSC. We injected AAV2-EF1a-DIO-hChR2(H134R)-eYFP into the NRM of *ePet-Cre* mice and implanted an optical fiber above the DHSC to directly stimulate the 5-HT fibers of the NRM projecting to the DHSC (5-HT^NRM^::hChR2 → DHSC mice, **Figure 1C**)^30,31^. A group of mice expressing only GFP into 5-HT^NRM^ was used as a control (5-HT^NRM^::GFP mice). We observed that stimulation at 5Hz/100ms induced a decrease in the PWT, promoting mechanical hypersensitivity mimicking the effect observed by chemogenetic activation of 5-HT^NRM^ cell bodies. In contrast, no effect was observed in control 5-HT^NRM^::eGFP mice (**Figure 1C-D**). This result was confirmed using repetitive subthreshold mechanical stimulations, which became suprathreshold during optogenetic stimulation (**SupFig3D**). We assessed thermal sensitivity using Hargreaves/Plantar test and found that optogenetic stimulation decreased the paw withdrawal latency, indicating thermal hypersensitivity (**SupFig3E**). Since both males and females were included in the study, we compared the effects of optogenetic stimulation and found mechanical hypersensitivity in both sexes (**SupFig3F-G**), suggesting no sex specific effect. Together, these data demonstrate that activation of the 5-HT^NRM^ → DHSC circuit increases sensitivity to nociceptive stimuli.

We then turned our attention to the effect of 5-HT^NRM^ manipulation within the DHSC on second order neuron activity. This was achieved using spinal cord *in vivo* electrophysiology in 5-HT^NRM^:hChR2 → DHSC mice. We determined the effect of 5-HT^NRM^ → DHSC stimulation on spinal excitability by recording DHSC neurons and their response to peripheral electrical stimulation of the ipsilateral paw which activates nociceptive fibers (**Figure 1E-G**). We found that optogenetic 5-HT^NRM^ → DHSC stimulation induced an increase in DHSC neuron excitability, as we observed a significant increase in the neuronal response to c-fiber stimulation, not present neither in GFP mice nor in fast non nociceptive A component (**Figure 1H and SupFig4**). In contrast, the amplitude of the windup in DHSC neurons, a marker of central sensitization, was not modified suggesting an upstream effect of DHSC maybe directly on c-fibers terminals (**Figure 1I-J**).

### Activation of 5-HT^NRM^ **→** DHSC neurons activation increases nociceptive sensitivity via 5-HT_3_ receptors

We next investigated the receptor target of 5-HT^NRM^ → DHSC projections involved in processing facilitation, focusing on 5-HT_3_ receptors, previously shown to mediate nociceptive facilitation in the DHSC^1–3^. 5-HT_3_ receptor is a ligand-gated ion channel that promotes neuronal depolarization and excitation. Using the RNAscope technique, we first visualized 5-HT_3_ receptor expression in the DHSC and found that its mRNA (*HTR3A*) is significantly more abundant in excitatory (glutamatergic) than in inhibitory (GABAergic) neurons (**Figure 2A-B**). After confirming the proalgesic effect of the optogenetic stimulation at 5Hz/100ms in 5-HT^NRM^:hChR2 → DHSC mice, we intrathecally injected the same animals with the 5-HT_3_ receptor antagonist granisetron (15μg). The optogenetically induced mechanical hypersensitivity in the 5-HT^NRM^ → DHSC circuit (**Figure 2C-D**) was completely abolished by 5-HT_3_ receptor blockade and a slight but significant analgesia was observed, suggesting that 5-HT_3_ receptor are involved in spinal mechanical hypersensitivity through activation of DHSC excitatory neurons (**Figure 2E**).

**Fig 2.**
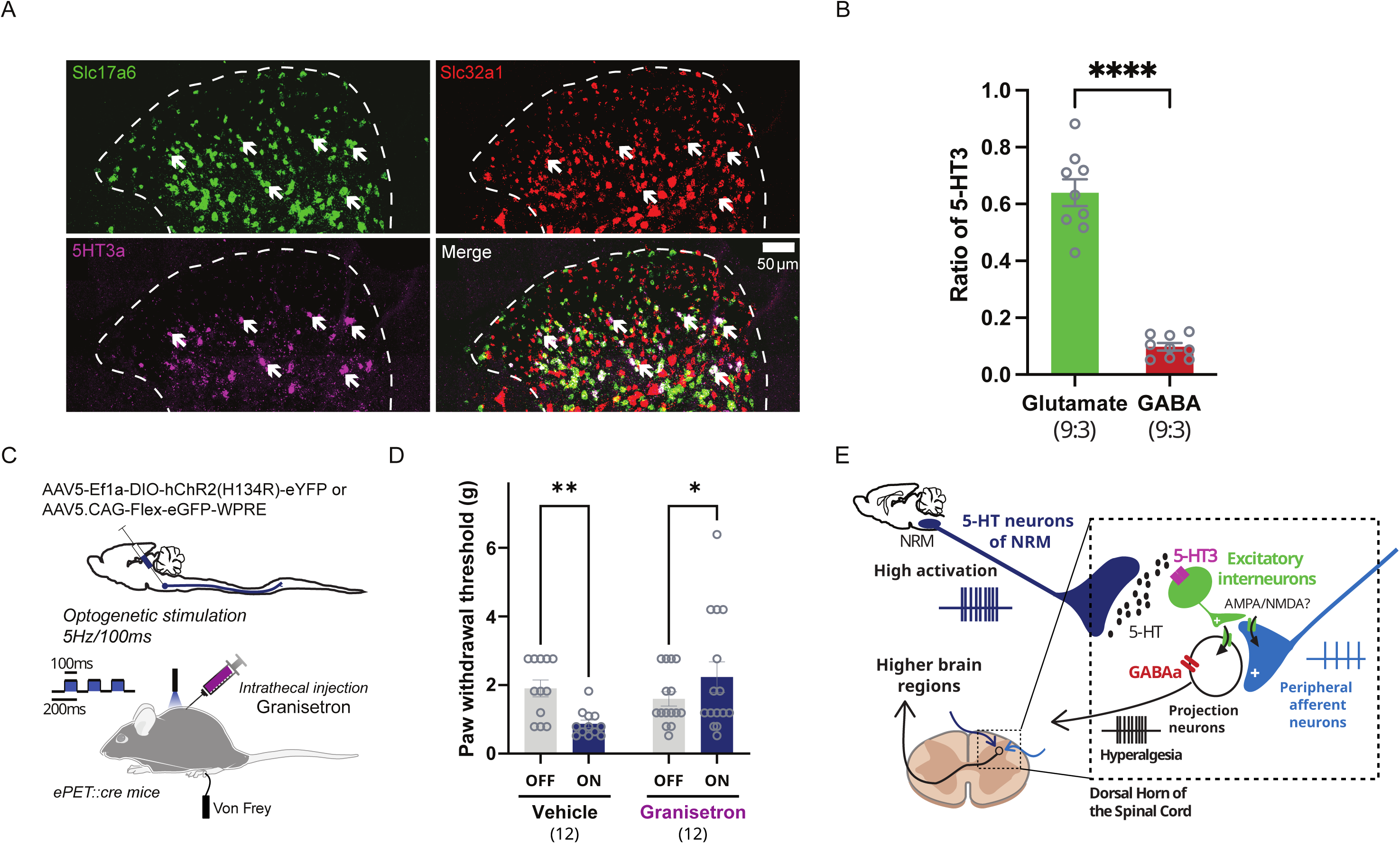
5-HT_3_ receptors mediate mechanical hypersensitivity through glutamatergic neurons. (A) Confocal images of DHSC of mice after RNAscope hybridization: 5-HT_3A_ receptors in magenta, Slc17a6 (glutamate) in green and Slc32a1 (GABA) in red. Colocalization of glutamate and 5-HT_3A_ is shown with white arrows. (B) In mice DHSC, 5-HT_3_ receptors are mostly colocalized with glutamatergic neurons (Unpaired t test, p<0.0001). Number in brackets (number of slices; number of animals). (C) Experimental protocol: NRM 5-HT descending fibers of ePet-Cre mice are stimulated optogenetically at 5Hz/100ms and the PWT of hinpaws is tested. (D) Optogenetic stimulation of 5-HT descending fibers after intrathecal injection of either vehicle (saline) or granisetron (Two-way ANOVA, p<0.0001). (E) Summary scheme of the putative mechanism of hyperalgesia induced by 5Hz/100ms stimulation of 5-HT^NRM^. Data are presented in mean ± SEM. Number in brackets = number of animals.

### inactivation of 5-HT^NRM^ neurons increases nociceptive sensitivity

In the next step, we aimed to determine whether 5-HT^NRM^ neurons are required to appropriately process nociceptive stimuli. We used the same approach as before by expressing the DREADD inhibitory receptor hM4(G_i_) exclusively in 5-HT^NRM^ neurons (**Figure 3A**). We then treated mice with saline (vehicle) or designer drug DCZ (0.1mg/kg, *i.p.*), which activates the hM4(Gi) receptor, and assessed nociception using the Von Frey test. Surprisingly, similar to activation, inhibition of 5-HT^NRM^ neurons by DCZ also decreased PWT (**Figure 3B**). This effect was reversible, as PWT returned to baseline 4 hours after DCZ injection. In contrast, no behavioral changes were observed in 5-HT^NRM^::hM4Di mice treated with vehicle or in control 5-HT^NRM^::eGFP mice treated with DCZ (**Figure 3B and SupFig5**). These results reveal that the basal activity of 5-HT^NRM^ neurons is crucial for the perception of nociceptive sensitivity, with both a decreased and an increased in 5-HT^NRM^ activity enhancing sensitivity, probably through different mechanisms.

**Fig 3.**
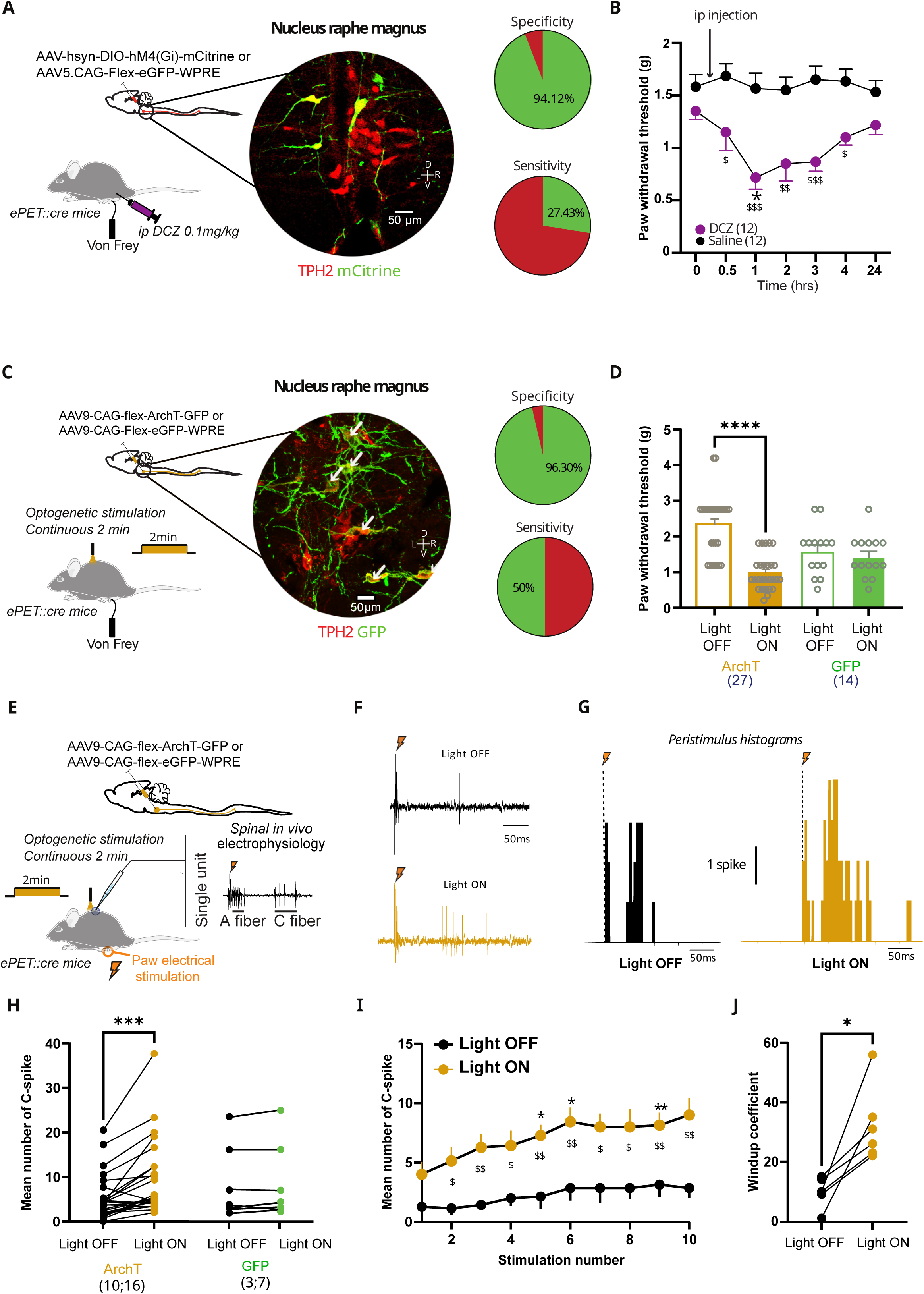
Inhibition of 5-HT neuron induces mechanical allodynia. (A) ePet-cre mice injected with a cre-dependent AAV to express the inhibitory DREADD hM4D. Expression of the mCitrine specifically colocalized with TPH2 immunostaining. (B) ip injection of DCZ induces a significant decrease in PWT that lasts for 3hrs but not with saline (Two-way ANOVA, p<0.01, * vs 0 and $ DCZ vs saline).(C) ePet-cre mice injected with a cre-dependent AAV to express the inhibitory opsin ArchT, optogenetic activation of ArchT is performed with an optic fiber placed above the lumbar spinal cord. Expression of the GFP specifically colocalized with TPH2 immunostaining. (D) Optogenetic inhibition of 5-HT fibers induces a significant decrease in PWT (Two-way ANOVA, p<0.0001). (E) ePet-cre mice injected with a cre-dependent AAV to express inhibitory opsin Archt in 5-HT neurons. (F) example of DHSC neuron response to supraliminar electric stimulation of the peripheral receptive field before (upper panel) and during optogenetic stimulation (lower panel). (G) peristimulus histogram of DHSC neurons before (left panel) and during (right panel) optogenetic inhibition of 5-HT neurons. (H) Number of c-spikes is significantly increased during optogenetic inhibition in ArchT but not in GFP group (Two-way ANOVA, p<0.05). Number in brackets (number of animals; number of neurons). (I) Windup curve is significantly increased during optogenetic inhibition of the 5-HT fibers (Two-way ANOVA, p_rank stim_<0.01, p_opto_<0.01). (J) Windup coefficient is significantly increased in mice expressing ArchT (Wilcoxon’s test,p<0.05). Data are presented in mean ± SEM.

To elucidate the circuit by which inhibition of 5-HT^NRM^ neurons increases pain processing, we used optogenetics. Control validation experiments were performed using patch clamp recordings on NRM brain slices. These studies confirmed that green light hyperpolarizes and inhibits ArchT-positive neurons in NRM slices of 5-HT^NRM^::ArchT mice (**SupFig6**). Then, we stereotactically injected the Cre-dependent virus AAV2.9-CAG-Flex-ArchT-GFP into the NRM of *ePet-Cre* mice, enabling the expression of the inhibitory opsin ArchT in 5-HT^NRM^ neurons (5-HT^NRM^::ArchT mice), or a control virus as previously described (5-HT^NRM^::eGFP mice) (**Figure 3C**). Then, in *ePet::cre* mice expressing the inhibitory opsin ArchT in the NRM, we optogenetically inhibited 5-HT projections from the NRM to the DHSC using continuous green light (5mW). Consistent with the chemogenetic inhibition of 5-HT neurons in the NRM, selective inhibition of 5-HT^NRM^::ArchT → DHSC also induced a significant decrease in PWT in 5-HT^NRM^::ArchT mice (**Figure 3D**). No such effect was observed in the 5-HT^NRM^::eGFP control mice. To confirm that inhibition generates hyperalgesia, we applied repetitive subthreshold stimulations to the paw using the Von Frey filament and observed that optogenetic inhibition induced an increase in the number of responses in 5-HT^NRM^::ArchT mice, an effect absent in 5-HT^NRM^::eGFP mice (**SupFig7B**). We tested for thermal sensitivity and found that optogenetic inhibition decreased the thermal latency indicating thermal hypersensitivity (**SupFig7C**). Both thermal and mechanical hypersensitivity induced by optogenetic inhibition were comparable between male and female mice (**SupFig7D-F**).

We then evaluated the effect of inhibition of descending 5-HT^NRM^ → DHSC fibers on the excitability of DHSC neurons. activation of nociceptive fiber was induced by electrical stimulation of the paw (**Figure 3E**). Optogenetic inhibition of 5-HT^NRM^ → DHSC fibers in 5-HT^NRM^::ArchT mice increased DHSC neuron excitability as evidenced by a significant increase in the number of c-spikes in response to electrical stimulation of the paw (c-fiber, **Figure 3F-H**). In contrast, no change was observed neither in 5-HT^NRM^::GFP mice nor in the A-fiber component (**Figure 3H and SupFig8**). Finally, we examined windup and found that optogenetic inhibition of descending 5-HT^NRM^ → DHSC fibers induced a substantial and significant increase in windup in 5-HT^NRM^::ArchT mice (**Figure 3I-J**). Together, these results show that inhibition of descending 5-HT pathway from the NRM to the DHSC promotes spinal hyperexcitability and nociceptive hypersensitivity, suggesting that 5-HT neurons exert a tonic analgesic effect under physiological conditions.

### 5-HT2C receptors mediates tonic spinal analgesia

We then examined which types of 5-HT receptors are mediating the analgesia induced by the modulation of 5-HT^NRM^ → DHSC pathway. Using publicly available datasets^21,22^, we found that the 5-HT_2C_ receptor subtype is highly enriched compared to the other fourteen 5-HT receptor subtypes in the DHSC. This enrichment is present in both excitatory and inhibitory subpopulations, regardless of the transcript extraction technique, whether single cell (**Figure 4A-B**) or single nucleus (**Figure 4C-D)**. Interestingly, the 5-HT_2C_ receptor is also enriched in human excitatory and inhibitory neurons based on single nuclei analysis of DHSC (**Figure 4E-F**). To confirm the presence of 5-HT_2C_ in mouse DHSC and their localization in inhibitory and excitatory neurons, we then used RNAscope and immunostaining techniques. We observed that endogenous *HTR2C* mRNA is expressed in both excitatory (glutamatergic) and inhibitory (GABAergic) neurons (**Figure 4G**) with a higher prevalence of *HTR2C* mRNA in excitatory neurons than inhibitory neurons (**Figure 4H**). This finding was confirmed in *5-HT_2C_-Cre* mice crossed with Ai9tdTomato reporter mice (*5-HT_2C_-Cre::Ai9* mice) to facilitate 5-HT_2C_ receptor visualization. Once again, we found that 5-HT_2C_ receptors are expressed in both excitatory (Tlx3-labeled) and inhibitory (Pax2-labeled) DHSC neurons, but in a greater number of excitatory neurons (**Figure 4I-J**). These results show a high level of expression of 5-HT_2C_ receptors in the DHSC in rodent and human with a higher prevalence in excitatory neurons but nevertheless a high expression in inhibitory neurons. We then tested the involvement of 5-HT_2C_ receptors in tonic analgesia by intrathecal injection of 5-HT_2_ antagonists (**Figure 4K**), i.e., methysergide, a 5-HT_2A-C_ and 5-HT_7_ receptor antagonist, ketanserin, a selective 5-HT_2A_ receptor antagonist and RS102221, a selective 5-HT_2C_ receptor antagonist. While RS102221 significantly reduced PWT, methysergide induced a slight but not significant decrease in PWT and ketanserin did not change it (**Figure 4L**). These results suggest that tonic 5-HT mechanical analgesia acts mainly via 5-HT_2C_ receptors.

**Fig 4.**
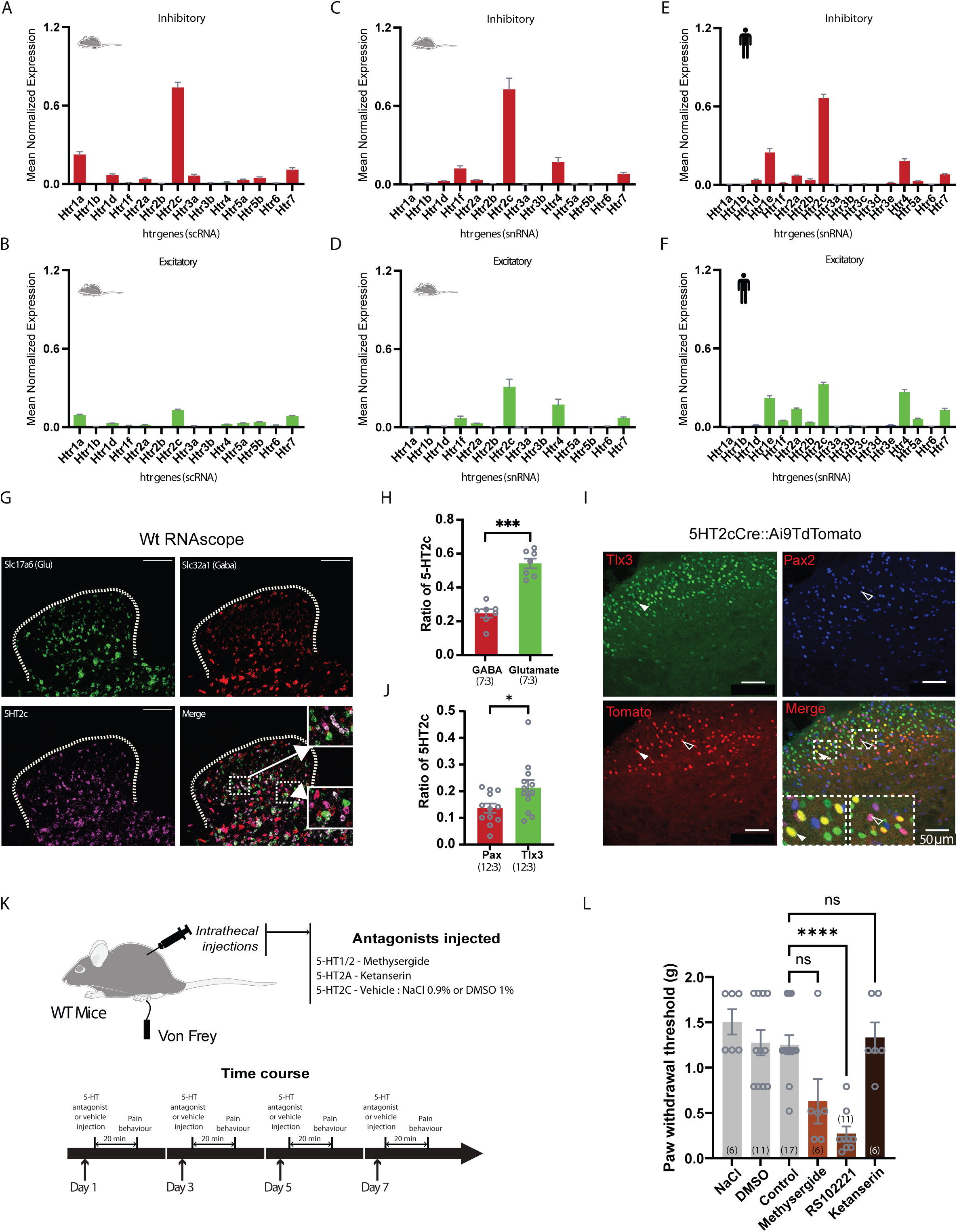
5-HT_2c_ receptors mediate mechanical analgesia. (A) Normalised expression of fourteen 5-HT receptors mRNA from single cell (sc) extraction (htr) in the inhibitory population of the mouse DHSC. (B) Normalised expression of fourteen 5-HT receptors mRNA (htr, sc) in the excitatory population of the mouse DHSC. (C) Normalised expression of fourteen 5-HT receptors mRNA from single nucleus (sn) extraction (htr) in the inhibitory population of the mouse DHSC. (D) Normalised expression of fourteen 5-HT receptors mRNA from sn extraction (htr) in the excitatory population of the mouse DHSC. (E) Normalised expression of fourteen 5-HT receptor mRNA from sn extraction (htr) in the inhibitory population of the human DHSC. (F) Normalised expression of fourteen 5-HT receptors mRNA from sn extraction (htr) in the excitatory population of the human DHSC. (G) Representative images of RNAscope for glutamate (Slc17a6), GABA (Slc32a1) and 5-HT_2c_ mRNA. Insets : higher magnification showing colocalization between glutamate and 5-HT_2c_ (upper panel) and between GABA/5-HT_2c_ (lower panel). (H) Quantification of colocalization shows 5-HT_2_c mRNA is present in both excitatory and inhibitory neurons but with a significant higher proportion in excitatory neurons (Paired t test, p<.001). (I) Confocal images of DHSC of 5-HT_2c_ Cre*Ai9 mice. A 5-HT_2c_ colocalization with Tlx3 is shown by the white arrow head while a colocalization with Pax2 is shown by the black arrow head. Insets: higher magnification. (J) In mice DHSC, 5-HT_2c_ are more colocalized with Tlx3 (glutamatergic neurons) than Pax2 (GABAergic neurons) (Paired t test, p<0.05). (K) Experimental design: PWTs of mice are tested after intrathecal injections of 5-HT2 antagonists. (L) Hyperalgesic effect of intrathecal injections of 5-HT_2c_ antagonists (Kruskal-Wallis, p<0.0001). Data are presented in mean ± SEM. Number in brackets =number of animals.

### 5Hz/5ms stimulation of 5-HT neurons induces analgesia

These results suggest that both inhibition and stimulation of the 5-HT^NRM^ → DHSC pathway are pronociceptive. To resolve this paradox, we modified the optogenetic stimulation parameters of the 5-HT^NRM^ → DHSC projection to determine whether the activation of 5-HT^NRM^ projecting DHSC neurons could be bidirectional depending on the activity of these neurons. To this end, we used a stimulation protocol corresponding to the normal activity level of 5-HT neurons^25,34^ (**Figure 5A**). This optogenetic stimulation protocol already has previously demonstrated a sex-independent mechanical and thermal analgesia ^25^. Here we confirmed this result as optogenetic stimulation of 5-HT^NRM^:hChR2 → DHSC with 5Hz/5ms, induced a significant increase in PWT not present in GFP mice (**Figure 5B**) and a decrease in the percentage of response to repetitive subthreshold stimulations (**SupFig9B**). In a separate group of mice, we tested the effect of both stimulation protocols on PWT and found an increase in PWT with the 5Hz/5ms protocol and a decrease in PWT with the 5Hz/100ms protocol (**SupFig9C-E)**.

**Fig. 5.**
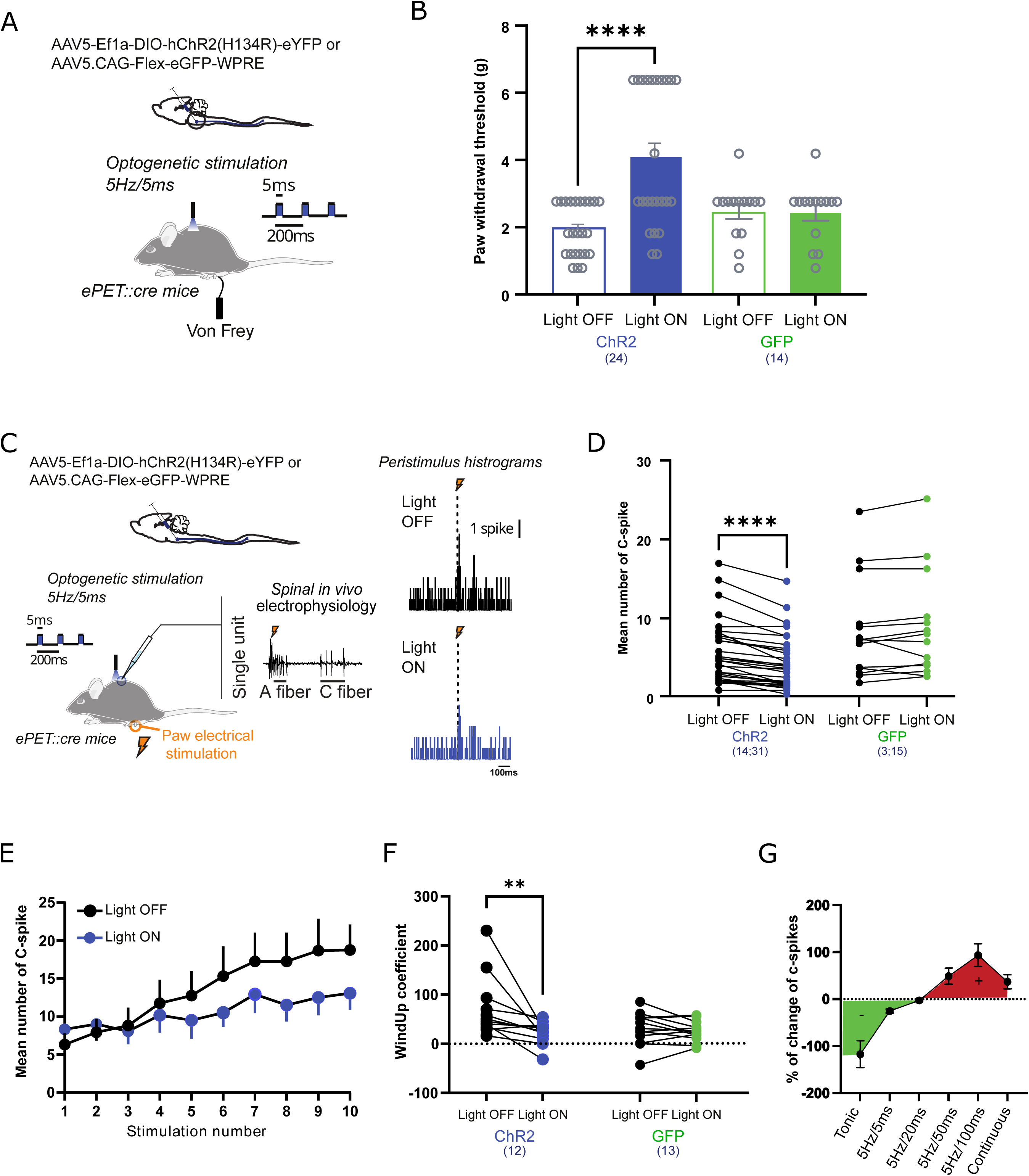
Optogenetic stimulation at 5Hz/5ms is analgesic. (A) experimental design as in fig 1C-D. (B) optogenetic stimulation of 5-HT fibers at 5Hz/5ms induces a significant increase in PWT (Two-way ANOVA, p<0.0001). (C) Extracellular recordings and peristimulus histogram of DHSC neurons before and during optogenetic stimulation at 5Hz/5ms. (D) The number of c-spikes is significantly decreased by low optogenetic stimulation in ChR2 but not in GFP mice (Two-way ANOVA, p<0.0001). Number in brackets (number of mice; number of neurons). (E) Windup is decreased by optogenetic stimulation at 5Hz/5ms (Two-way ANOVA p_opto_>0.05, p_rank of stim_<0.0001). (F) windup coefficient is significantly decreased in Chr2 group but not in GFP (Two-way ANOVA, p<0.05). Number in brackets= number of animals. (H) Modulation of 5-HT neurons and change in DHSC neurons response. Data are presented in mean ± SEM.

Next, we performed *in vivo* electrophysiological recordings from DHSC neurons in anesthetized 5-HT^NRM^:hChR2 → DHSC mice (**Figure 5C**). We observed that optogenetic 5Hz/5ms 5-HT^NRM^ → DHSC stimulation induced a reduction in the c-fiber response of DHSC neurons in response to peripheral nociceptive stimuli but not in GFP mice (**Figure 5D**). We observed no change in the A component (**SupFig9F-H**). The windup coefficient was significantly decreased, suggesting a direct action of 5-Hz/5ms optogenetic stimulation on DHSC neurons in 5-HT^NRM^::hChR2 → DHSC mice (**Figure 5E-F**). 5-HT^NRM^:hChR2 → DHSC exerts a bidirectional control of DHSC neuron excitability, including inhibition-induced hyperexcitability, short-stimulation-induced decreased excitability and long-stimulation-induced hyperexcitability (**Figure 5G**). Finally, to confirm that this difference is due to the level of activation and not to the recruitment of different 5-HT neuronal subpopulations, we performed single whole-cell recordings of 5-HT neurons of the NRM using *5-HT::Ai9 tdtomato* mice. By comparing their somatic electrophysiological properties, we found that 5-HT^NRM^ neurons represent a homogenous population with similar intrinsic properties (**SupFig10**).

### 5Hz/5ms optogenetic stimulation activates spinal inhibitory interneurons

In the next step, we aimed to identify the neurons receiving signals from 5-HT^NRM^:hChR2 → DHSC. First, under anesthesia, we stimulated 5-HT^NRM^ → DHSC at 5Hz/5ms for 10 minutes in 5-HT^NRM^:hChR2 → DHSC mice and measured the expression of the activity marker c-Fos in the DHSC *ex vivo* (**SupFig11A**). We found that the number of c-Fos*-*expressing neurons in the DHSC is significantly increased after optogenetic stimulation at 5Hz/5ms of 5-HT^NRM →^ DHSC pathway (**SupFig 11B**), and that this increase mainly involved neurons expressing Pax2, a marker of inhibitory neurons in the DHSC (**SupFig 11C**). Inhibitory neuronal subpopulations of the DHSC express several 5-HT receptors, including the G_i_-coupled inhibitory 5-HT_1_ receptor, the G_q_-coupled excitatory 5-HT_2_, 5-HT_4_ and 5-HT_7_ receptors, and the ionotropic 5-HT_3_ receptor (**Figure 4A, C, E**). Since c-Fos expression is induced in inhibitory neurons by optogenetic activation of the 5-HT^NRM^:hChR2 → DHSC, we excluded 5-HT_1_ receptors associated with inhibitory signaling. Among the excitatory 5-HT receptor subtypes, the most highly expressed was the 5-HT_2C_ receptor, but also with lower levels, the 5-HT_2A_, 5-HT_3A_, 5-HT_4_, and 5-HT_7_ receptors. Here, we focused on 5-HT_2A_, 5-HT_2C_ and 5-HT_3_ receptors. We have previously identified 5-HT_2C_ and 5-HT_3_ receptors in inhibitory neurons (**see Figure 2 and 4**), we assessed the expression of 5-HT_2A_ receptors. RNAscope analysis confirmed the presence of *HTR2A* mRNA in both excitatory and inhibitory neurons in the DHSC with a higher prevalence in inhibitory neurons (**SupFig 11D1-D2**). The results were confirmed using the *5-HT_2A_R-Cre* transgenic mice crossed with an Ai9-Tdtomato reporter mice where we found that 5-HT_2A_ receptors are almost exclusively expressed in inhibitory neurons (**SupFig 11E1-E2).**

### 5Hz/5ms-induced analgesia is mediated by 5-HT_2A_ and 5-HT_2c_ receptors

Among the three receptors identified in inhibitory neurons (5-HT_2A_, 5-HT_2C_ and 5-HT_3_), we tested the one responsible for the analgesia induced by optogenetic stimulation at 5Hz/5ms of 5-HT^NRM^ → DHSC neurons. We injected intrathecally 5-HT_2A_, 5-HT_2C_ or 5-HT_3_ receptor antagonists into 5-HT^NRM^::hChR2 → DHSC mice and evaluated the analgesic effect of 5Hz/5ms stimulation of descending 5-HT fibers on the PWT. After methysergide injection, optogenetic stimulation of 5-HT^NRM^:hChR2 DHSC pathway no longer had an analgesic effect (**Figure 6A**). The same result was observed after injection of RS102221 and ketanserin (**Figure 6B-C**). In contrast, the 5-HT_3_ receptor antagonist granisetron did not modify the analgesic effect of optogenetic stimulation of 5-HT^NRM^:hChR2 → DHSC (**Figure 6D**), in line with the low expression of 5-HT_3_ receptors in inhibitory neurons of the DHSC. The absence of effect of granisetron rules out the involvement of 5-HT_3_ receptors, but the similar action of methysergide, ketanserin and RS10221 supports the hypothesis of 5-HT_2C_/5-HT_2A_ receptor-dependent analgesia for 5Hz/5ms stimulation (**Figure 6E-F**).

**Fig. 6.**
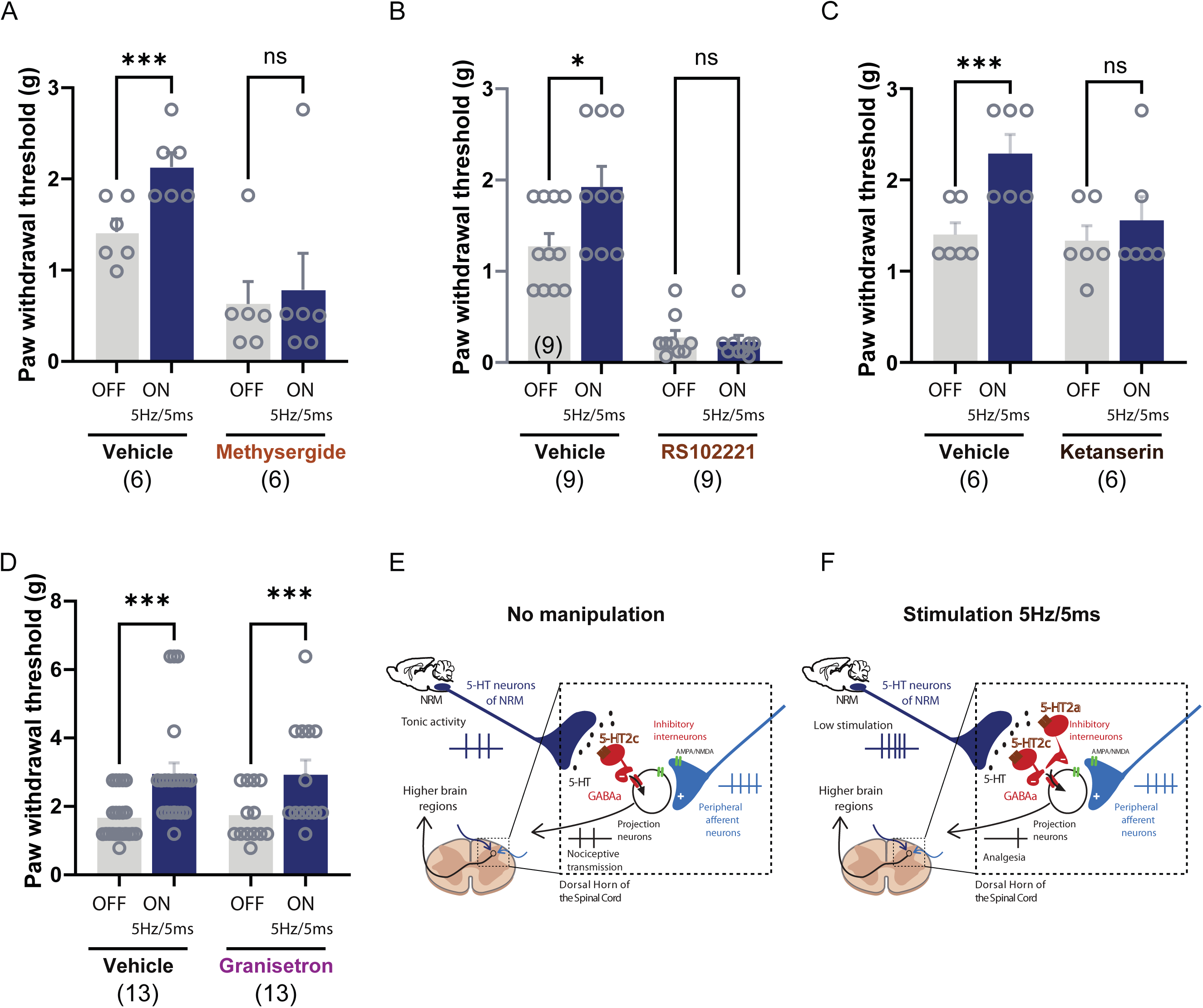
Analgesic effect of optogenetic stimulation of 5-HT fibers is mediated by 5-HT_2c_ and 5-HT_2A_ receptors. (A) Optogenetic stimulation of 5-HT fibers significantly increased PWT after intrathecal saline but not after intrathecal injection of the 5-HT_2_ antagonist methysergide (Two-way ANOVA, p<0.01). (B) Optogenetic stimulation of 5-HT fibers significantly increases PWT after intrathecal saline but not after intrathecal injection of 5-HT_2c_ antagonist RS10221 (Two-way ANOVA p<0.05). (C) Optogenetic stimulation of 5-HT fibers significantly increased PWT after intrathecal saline but not after intrathecal injection of 5-HT_2A_ antagonist ketanserin (Two-way ANOVA, p<0.001). D) Optogenetic stimulation of 5-HT fibers significantly increases PWT after intrathecal saline or 5-HT_3_ antagonist granisetron (Two-way ANOVA, p>0.05) (E) Mechanism of tonic spinal analgesia by 5-HT^NRM^ neurons. (E) Mechanism of spinal analgesia induced by 5Hz/5ms activation of 5-HT^NRM^ neuron. Data are presented in mean ± SEM. Number in brackets define the number of animals.

These results provide evidence for a model in which 5-HT^NRM^ neurons projecting to the DHSC serve as regulators to amplify or attenuate pain-related stimuli depending on the level of 5-HT released in DHSC (**SupFig12).**

## DISCUSSION

In the present study, we aimed to define the role of the activity level of 5-HT^NRM^ neurons in the brainstem on nociceptive transmission and spinal excitability. We show that this population of 5-HT neurons, connected to the spinal inhibitory network, exerts a tonic inhibitory effect that transforms into excitation at a higher activity level. More specifically, under physiological conditions, 5-HT^NRM^ neurons projecting to the DHSC exert a tonic analgesic effect through 5-HT_2C_ receptors. We also show that a slight increase in the activity of fibers originating from 5-HT^NRM^ and projecting to the DHSC enhances the analgesic effect by recruiting 5-HT_2A_ receptors, while more intense stimulation induces hyperalgesia via 5-HT_3_ receptors.

The effect of 5-HT application/injection on nociception and spinal excitability has been studied for a long time, and results in both pro- and antinociceptive action^10,11,35^. Bath application of 5-HT on acute spinal slices, induces both an outward and an inward current depending on the recorded neuron of the *substantia gelatinosa*, an important region of nociceptive control^16^. Particularly, vertical cells, known as excitatory interneurons respond to 5-HT with an outward current suggesting a decrease in neuronal excitability while 5-HT induces both an inward and an outward current on islet cells, inhibitory interneurons suggesting dual action of 5-HT on this neuronal population^16,36^. 5-HT depresses the sensory transmission by decreasing excitatory postsynaptic potentials in most of intracellularly recorded spinal neurons but also opposite actions in a subset of these neurons^14,15^. 5-HT also decreases the c-fiber response of extracellularly recorded wide dynamic range DHSC neurons^37^. At the behavioral level, 5-HT has also a bidirectional effect depending on the concentration used^17^. It is very likely that the effect of 5-HT on DHSC neurons depends on the dose used and the type of neuron recorded^38^. Here, by manipulating the 5-HT^NRM^, we confirm this bidirectional effect of spinal 5-HT and demonstrate that this effect is reproduced by 5-HT coming from brainstem 5-HT nuclei.

Since the DHSC does not contain 5-HT neurons, most of the actions of 5-HT in the spinal cord originate from descending pathways in the brainstem, including the NRM. The consequences of electrical stimulation of the NRM on spinal transmission have been studied in various animal models from turtles to cats^39–41^. This stimulation induces both antinociception and a decrease in nociceptive transmission in the spinal cord^39,42–46^. Interestingly, the strongest inhibition is achieved with a lower stimulation intensity in the NRM ^47^. This antinociceptive effect likely involves brainstem 5-HT neurons, as electrical stimulation of the descending pathways, particularly the periaqueductal gray (PAG) and NRM, induces 5-HT release in the DHSC^44,48–53^. Taken together, these data suggest that 5-HT^NRM^ neurons induce spinal 5-HT release, thereby modulating nociceptive transmission. However, electrical stimulation can activate many different neuronal subpopulations in the NRM and spinal 5-HT could originate from different regions of the midbrain and brainstem. In a recent study from our laboratory, we showed that specific optogenetic activation of 5-HT^NRM^ induces increase 5-HT levels in the DHSC^25^. In the present study, we first show that chemogenetic activation of the 5-HT^NRM^ induces a pronociceptive response, a result similar to that of previous studies^23,24^. We reproduced this pronociceptive effect using long-duration (100 ms) repetitive optogenetic stimulation (5 Hz) and obtained an effect similar to the previous results with high frequency stimulation (25Hz)^23^. Moreover, by stimulating selectively the 5-HT^NRM^-DHSC pathway, we show that this effect is directly due to the projections of 5-HT neurons to the spinal cord. This result undoubtedly confirms that 5-HT neurons in the NRM induce an increase in nociceptive response via a direct 5-HT^NRM^-DHSC pathway when they are overactivated beyond their tonic discharge (<5Hz)^34^.

Whether 5-HT^NRM^ neurons exerts a tonic descending control of nociception has long been debated, and the role of this tonic activity remains poorly defined^45,54,55^. Our results clearly show that 5-HT^NRM^-DHSC projections exert tonic inhibition on DHSC neurons and tonically attenuate nociception. This tonic action of brainstem 5-HT neurons has also been demonstrated in pathological pain, with an inhibitory or facilitatory role depending on the cause of the pain hypersensitivity. Indeed, in the model of neuropathy induced by peripheral nerve injury, 5-HT^NRM^ are pronociceptive and increase the excitability of DHSC neurons^25,56,57^. The shift from antinociception to pronociception could be due to a loss of inhibitory controls associated with chloride imbalance in the DHSC^25,58,59^. Conversely, 5-HT^NRM^ are antinociceptive in case of persistent inflammation or hypersensitivity associated with neurodegenerative disease^29,60^.

Short, low frequency stimulation activates spinal inhibitory interneurons, as it induces an increase in the number of c-Fos positive neurons that express the inhibitory marker Pax2. We have previously shown that intrathecal injection of picrotoxin, which blocks GABA_A_ receptors, suppresses the antinociceptive action of optogenetic stimulation of 5-HT^NRM^ ^25^. This result is consistent with previous studies reporting that 5-HT acts on DHSC inhibitory interneurons in the DHSC. Indeed, 5-HT increases the amplitude and frequency of inhibitory post-synaptic currents in DHSC neurons of the *substantia gelatinosa*. The frequency change under TTX, suggests a presynaptic action likely due to 5-HT receptors on inhibitory neurons^16^. Finally, in GAD67::GFP mice, 5-HT has been shown to induce an inward current in GAD(+) inhibitory interneurons associated with an increase in the frequency of miniature IPSCs in GAD(-) neurons suggesting that 5-HT activates inhibitory interneurons, thereby increasing the efficiency of inhibition on excitatory neurons^61^. We and others have shown that 5-HT^NRM^ do not project to the superficial lamina^25,57^. Given that the antinociceptive action of the optogenetic stimulation of 5-HT^NRM^ modifies both mechanical and thermal sensitivity, induces c-Fos expression in inhibitory interneurons in the deep DHSC and affects the excitability of wide dynamic range neurons, we propose that the spinal microcircuit involved is primarily located in the deep laminae of the DHSC. By contrast, longer and higher-frequency stimulation induces an increase in nociception. We hypothesize that this effect increases 5-HT release in the DHSC, which propagates into the spinal microcircuit and act via volumetric mechanisms.

At least 14 5-HT receptors are expressed in the DHSC^10,11,35^. Analyses of RNAseq datasets^62,63^ revealed that inhibitory interneurons express mainly 5-HT_1_, 5-HT_2_, and 5-HT_3_ receptors^21,22^. Since 5-HT_1_ receptors are coupled to Gi protein, the receptors most likely to activate inhibitory interneurons would be 5-HT_2_ or 5-HT_3_ receptors. Using transgenic mice and the RNAscope technique, we show that among these receptors, 5-HT_2A/C_ receptors are more highly expressed in inhibitory interneurons than 5-HT_3_ receptors. We show that the tonic antinociceptive effect of 5-HT^NRM^ is mediated by 5-HT_2c_ receptors and that the antinociception induced by brief stimulation of 5-HT^NRM^ is mediated by both 5-HT_2A_ and 5-HT_2c_ receptors. This effect is likely due to the excitatory role of both 5-HT_2_ receptors located on inhibitory interneurons, as this stimulation protocol activates these interneurons. The role of 5-HT_2c_ receptors in spinal nociceptive transmission has been underestimated and only few studies suggest a role of these receptors in nociception although their role on other physiological processes is well established (e.g., anxiety, stress-induced food intake ^64–66^). In the case of pathological pain, some studies have highlighted the possible role of 5-HT_2c_ receptor agonists in relieving pain hypersensitivity, with these modulators acting either centrally or peripherally^67,68^. Here, using a large open access RNAseq dataset, we show that 5-HT_2c_ receptors are the most highly expressed 5-HT receptor in the DHSC of rodents, as well as in human tissue. Furthermore, in both rodents and humans, 5-HT_2c_ receptors are highly expressed in inhibitory interneurons, reinforcing the translatability of our findings to humans. The use of RNAscope technology which improves sensitivity, confirms the expression of 5-HT_2c_ receptors in both inhibitory and excitatory neurons, with slightly higher expression in the latter. The role of 5-HT_2c_ receptors in excitatory neurons will need to be further investigated before developing new analgesic compounds targeting these receptors.

We demonstrate that antinociception induced by brief optogenetic stimulation also recruits 5-HT_2A_ receptors. Previous studies on the role 5-HT_2A_ receptors in pain and nociceptive transmission show that these receptors are involved in nociceptive transmission and that their action, anti- or pronociceptive depends on their location, peripheral or central, and on the nature of the neurons, excitatory or inhibitory. In the DHSC, 5-HT_2A_ receptor agonists reduce the excitability of DHSC neurons and this 5-HT inhibitory effect is blocked by 5-HT_2A_ receptor antagonists ^37^. This is consistent with findings showing that intrathecal injection of 5-HT_2_ receptor agonists have an anti-allodynic effect in a model of peripheral neuropathy and that antagonists of these receptors have antinociceptive action in models of neuropathic pain ^69^. Here, we show that blocking the receptor located on inhibitory interneurons abolished the antinociceptive effect. The antinociceptive role of 5-HT_2A_ receptor is an important translational result as recent years have seen a resurgence of interest in psychedelic molecules for the treatment of neurological disorders, including pain, and these hallucinogenic and non-hallucinogenic molecules target the 5-HT_2A_ receptors^70–72^.

The pronociceptive role of 5-HT_3_ receptors has long been demonstrated, and the use of 5-HT_3_ receptor antagonists has been proposed as pain killer. 5-HT_3_ receptors have been shown to be involved in the descending facilitation of nociceptive transmission mediated by 5-HT^73–75^. However, despite promising preliminary results from case studies, 5-HT_3_ receptor antagonists are still not recommended for the treatment of chronic pain^21,76^. Here, we show that prolonged optogenetic or chemogenetic stimulation of 5-HT^NRM^ induces pronociception, which is suppressed by granisetron, a 5-HT_3_ receptor antagonist. Analysis of RNAscope data showing the presence of 5-HT_3_ receptor mRNA mainly in excitatory neurons, strongly suggests that 5-HT acts on 5-HT_3_ receptors in these neurons. The lack of proof of concept for the efficacy of 5-HT_3_ receptor antagonists in humans may be explained by the fact that, in some publications, 5-HT_3_ receptors also possess an antinociceptive effect. Indeed, 5-HT_3_ receptors can depolarize inhibitory neurons, increase inhibitory postsynaptic currents on excitatory neurons, and decrease the excitability of DHSC neurons^37,61,77^. Furthermore, analysis of RNAseq datasets from human DHSC indicated very low expression of 5-HT_3_ receptors which could explain the lack of transferability of preclinical findings regarding this receptor. Some studies have shown that 5-HT_3_ receptors could act through sensory fibers^78^, but again data from human dorsal root ganglia found very low expression of 5-HT_3_ receptor mRNA in dorsal root ganglion neurons^79^.

Overall, our results demonstrate that descending 5-HT^NRM^ neurons projecting to the DHSC modulate the amplification or attenuation of painful stimuli depending on the degree of 5-HT released into the DHSC. More specifically, within physiological limits, increased activity of 5-HT neurons induces analgesia, while high activity induces pain hypersensitivity. It is noteworthy that these two distinct effects are controlled by different 5-HT receptors located on different spinal cord microcircuit, with or without inhibitory interneurons: 5-HT_2_ receptors inhibit pain, and 5-HT_3_ receptors promote hyperalgesia. Finally, RNAseq data from the human DHSC suggest that 5-HT_2C_ receptors represent a promising target for future exploration.

## Acknowledgements

We would like to thank Antoine Godin (Cervo) for kind help and advices concerning immunolabelling. We thank Dr. Thomas Müller for providing Tlx3 antibodies. We also thank the Bordeaux Bordeaux Imaging Center, BIC, UMS, CNRS, Université de Bordeaux and the Histocare platform, IMN CNRS UMR5293.

## Author Contribution

Z.G., A.V., P.F., Ab.B., YdK conceived and designed the experiments; Z.G., A.V., F.A., R.B.B., F.N., H.M., T.D., M.B., E.P. and A.R. collected data; F.M, Z.G., A.V., X.F., M.B., performed analysis; T.D., M.L. and A.B., developed tool analysis; An.B. L.K.H. prepared experimental transgenic mice; Ab.B., Z.G., P.F., wrote the paper; An.B, A.V., R.B.B., YdK, X.F. and L.K.H. carefully read and edited the manuscript.

