## Supplementary material for "Nucleus raphe magnus serotonin neurons bidirectionally control spinal nociceptive transmission in mice": Sup Fig Legend

**Supplementary materials**

*Sup Fig 1: Experimental designs. (A) Chemogenetic. (B) Optogenetic. (C) Optogenetic and pharmacology.*

*Sup Fig 2: Micrographs of the brain area infected after virus injection and related atlas (AAV5-Ef1a-Dio-hChR2(H134R)-eYFP) in one ePet-cre mice (scale: 1mm). Left rostral. right caudal. Virus infection in the nucleus raphe magnus (scale : 100μm). Bottom right: graph showing the number of cells infected according to the distance from the bregma (Three slices from one mice). Data are presented in mean ± SEM.*

*Sup Fig 3: (A) In mice injected with either hM3(Gq) or GFP : (B) time course of action of ip DCZ injection on the paw withdrawal threshold. DCZ has no effect on GFP mice while significantly decreasing the PWT in hM3(Gq) group (Two-way ANOVA, p<0.001). (C) Mice infected with either ChR2 or GFP expressing viruses. (D) repetitive mechanical subthreshold stimulation induced suprathreshold response during optogenetic 5Hz/100ms stimulation of 5-HT^NRM^ neurons. This effect is not present in GFP mice (Two-way ANOVA, p<0.0001). (E) Optogenetic stimulation at 5Hz/100ms induces a significant decrease in thermal latency in ChR2 group of mice but not in GFP mice (Two-way ANOVA, p<0.001). (F-G) Comparison of optogenetic stimulation of 5-HT^NRM^ neurons at 5Hz/100ms in males and females. (F) Optogenetic stimulation induces significant decrease in paw withdrawal threshold in both males and females (Wilcoxon test, p_males_<0.001, p_females_<0.01). (G) Optogenetic stimulation induces significant increase in the number of responses in both males and females (Wilcoxon test, p_males_<0.05, p_females_<0.05). Data are presented in mean ± SEM.*

*Sup Fig 4: (A) experimental paradigm. (B) optogenetic stimulation at 5Hz/100ms does not change the number of a-spike of dorsal horn neurons in ChR2 and in GFP group (Wilcoxon test, p>0.05) (Number of mice; Number of neurons). Data are presented in mean ± SEM.*

*Sup Fig 5: (A-B) DCZ ip injection in mice expressing the inhibitory DREADD hM4(Gi) induces a significant decrease in PWT that is not present in GFP control group (Two-way ANOVA, p<0.05). Data are presented in mean ± SEM.*

*Sup Fig 6: Patch-clamp recordings of 5-HT neurons expressing the inhibitory opsin ArchT. (A) Image of identified neurons for patch clamp recordings. (B) In neurons expressing the opsin, spontaneous neuronal discharge is stopped by green light (Opto). It generates a fast hyperpolarization followed by a long-lasting inhibition of the neuron. (C) Spontaneous activity is significantly decreased after continuous light (Paired t test, p<0.0001). Data are presented in mean ± SEM.*

*Sup Fig 7: (A) Mice infected with either ArchT or GFP expressing viruses. (B) Repetitive mechanical subthreshold stimulation induced suprathreshold response during optogenetic inhibition of 5-HT^NRM^ neurons. This effect is not present in GFP mice (Two-way ANOVA, p<0.0001). (C) Optogenetic inhibition of 5-HT^NRM^ neuron induces a significant decrease in paw withdrawal latency following thermal stimulation. This effect was not present in GFP mice. (Two-way ANOVA, p<0.0001). (D-F) Comparison of the effect of optogenetic inhibition of 5-HT^NRM^ neurons in males and females following mechanical or thermal stimulations. (D) Optogenetic inhibition induces significant decrease in paw withdrawal threshold in both males and females (Wilcoxon test, p_males_<0.05, p_females_=0.0001). (E) Optogenetic inhibition induces significant increase in the number of responses to subthreshold mechanical stimulations in both males and females (Wilcoxon test, p_males_<0.05, p_females_<0.0001). (F) Optogenetic inhibition induces significant decrease in thermal latency in both males and females (Paired t test, p_males_<0.01, p_females_<0.0001). Data are presented in mean ± SEM.*

*Sup Fig 8: (A) experimental paradigm. (B) optogenetic inhibition with ArchT illumination does not change the number of A-spikes in dorsal neurons after peripheral nociceptive stimulation (Paired t test, p>0.05) (Number of mice: Number of neurons). Data are presented in mean ± SEM.*

*Sup Fig 9: (A) experimental paradigm. (B) Repetitive mechanical suprathreshold stimulations are shifted to subthreshold during optogenetic 5Hz/5ms stimulation of 5-HT^NRM^ neurons . This effect is not present in GFP. (Two-way ANOVA, p<0.0001). (C-E) Effect of 5Hz/5ms and 5hz/100ms optogenetic stimulation in the same group of mice. (C) experimental paradigm. (D) 5Hz/5ms (left histogram) stimulation induces an increase in paw withdrawal threshold and 5Hz/100ms (right histogram) induces a decrease in paw withdrawal threshold (Two-way ANOVA, p<0.0001). (E) 5Hz/5ms (left histogram) stimulation induces an increase in paw withdrawal latency to thermal stimulation and 5Hz/100ms (right histogram) induces a decrease in paw withdrawal latency (Two-way ANOVA, p<0.0001). (F) experimental paradigm. (G-H) Optogenetic stimulation at 5Hz/5ms does not change the number of A-spikes in dorsal neurons after peripheral nociceptive stimulation in ChR2 nor GFP animals (Paired t test, p_ChR2_>0.05, p_GFP_>0.05). Data are presented in mean ± SEM.*

*Sup Fig 10: Electrophysiological properties and excitability of 5-HT neurons of the NRM in mice. (A) Representative traces of membrane potential of 5-HT neurons in response to either negative (-40 to -80pA) or positive (40 to 80pA) current injection. (B) Frequency distribution of the resting membrane potential of 5-HT neurons (n=23. binning 2 mV). (C) Frequency distribution of the membrane resistance of 5-HT neurons (n=23. binning 75MOhms). (D) Summary bar graph illustrating the membrane capacitance of 5-HT neurons (tau value 31.48±1.36ms. n=23). (E) Frequency distribution of the action potential threshold of 5-HT neurons (n=23. binning 2mV). (F) Frequency distribution of the rheobase of 5-HT neurons (n=23. binning 5 pA). (G) Summary bar graph illustrating the number of action potential upon the first supraliminal stimulation of 5-HT neurons (3.48±0.27. n=23). (H) Summary bar graph illustrating action potential time to peak of 5-HT neurons (1.06±0.04ms. n=23). (I) Summary bar graph illustrating action potential amplitude of 5-HT neurons (61.74±2.63 mV. n=23).*

*Sup Fig 11. 5Hz/5ms stimulations of 5-HT descending fibers activate inhibitory interneurons.*

*(A) c-Fos expression in mice DHSC after a 5Hz/5ms optogenetic stimulation of 5-HT fibers. Insets: higher magnification of Pax2/c-Fos colocalizations in superficial and deep laminae. (B) Increase of c-Fos positive cells after optogenetic stimulation of 5-HT fibers (Kruskal Wallis test, p<0.01). (C) This increase is present on inhibitory neurons (Kruskal Wallis test, p<0.01). (D1) RNAscope for Glutamate (Slc17a6), GABA (Slc32a1) and 5-HT2a mRNAs. (D2) quantification of colocalization between glutamate and 5-HT2a or Gaba and 5-HT2a mRNAs (Paired t test, p<0.0001). (E1) Confocal images of DHSC of 5-HT2aCre*Ai9 mice. A 5-HT2a colocalization with Pax2 is shown by white arrows. Inset: higher magnification. (E2) In mice DHSC, 5-HT2_A_Rs are more colocalized with Tlx3 (glutamatergic neurons) than Pax2 (GABAergic neurons) (Paired t test, p<0.0001). Data are presented in mean ± SEM. Number in brackets (number of slices:number of animals).*

*Sup Fig 12: General diagram of the activity patterns of 5-HT^NRM^ inducing analgesia or hyperalgesia.*
