## Supplementary material for "Nucleus raphe magnus serotonin neurons bidirectionally control spinal nociceptive transmission in mice": Table S4

| Figure N° | Mean ± SEM | Normality test  Shapiro-wilk test | Statistic method | Main effect | Post hoc | Number of mice (N) | Number of samples  (n) |
| --- | --- | --- | --- | --- | --- | --- | --- |
| S3B | **Gq DCZ**  baseline 1.367±0.02305  0.5Hr 1.017±0.06907  1Hr  0.8±0.06138  2Hr  0.9±0.06019  3Hr 1.217±0.07114  4Hr  1.2±0.06370  24Hr 1.517±0.06246  **GFP DCZ**  baseline 1.33±0.045  0.5Hr 1.28±0.162  1Hr 1.22 ±0.156  2Hr 1.2 ±0.15  3Hrs 1.13± 0.089  4Hrs 1.17 ±0.06  24Hrs 1.23 ±0.06 | No | Two Way RM ANOVA | p_interaction_=0.0005  F_interaction_=6.467 | Sidak’s  p_gq dcz vs gfp dcz_ ; _row3_ p=0.0261.  DCZ Gq  Baseline vs 0.5Hr p=0.0047 ; 1Hr p<0.0001 ; 2hrs p<0.0001.  DCZ GFP  p>0.05 | 24 Gq  12 GFP | NN |
| S3D | **ChR2**  Light OFF : 17.5%±3.9  Light ON  88.3%±3.7  **GFP**  Light OFF vs 10%±5.3  Light ON  10%±6.5 | No | Two Way RM ANOVA | p_interaction_<0.0001  F_interaction_=117.4 | Sidak’s  Light OFF vs Light ON ChR2 p<0.0001  Light OFF vs Light ON GFP p=0.999 | 12 ChR2  8 GFP | NN |
| S3E | **ChR2**  light ON  2.8±0.11s  light OFF.  4±0.08s  **GFP**  light ON  3.4±0.34s  light OFF  3.3±0.27s | No | Two Way RM ANOVA | p_interaction_<0.0006  F_interaction_=29.8 | Sidak’s  Light OFF vs Light ON ChR2 p=0.0002  Light OFF vs Light ON GFP p=0.991 | 5 ChR2  5 GFP | NN |
| S3F  Males | **ChR2**  light OFF. 1.6±0.17g  light ON  0.67±0.08g | No | Wilcoxon | p<0.001 | NN | 11 | NN |
| S3F  Females | **ChR2**  light OFF.  1.1±0.19g  light ON  0.5±0.1g | No | Wilcoxon | p=0.0078 | NN | 8 | NN |
| S3G  Males | **ChR2**  Light OFF  8±8%;  Light ON  92±4.9% vs | No | Wilcoxon | p=0.0312 | NN | 6 | NN |
| S3G  Females | **ChR2**  Light OFF  0±0%  Light ON  90±4.4% | No | Wilcoxon | p=0.0312 | NN | 6 | NN |
| S4B  ChR2 | **ChR2**  Light OFF  4±0.6  Light ON  4±0.6 | No | Wilcoxon | p=0.71 | NN | 8 | 25 |
| S4B  GFP | **GFP**  Light OFF  5.4±0.92  Light ON  5.7±1.11 | No | Wilcoxon | p=0.78 | NN | 4 | 17 |
| S5B | **Gi**  **DCZ**  Baseline 1.35±0.078  0.5Hr 1.15±0.176  1Hr 0.7167±0.1114  2Hrs 0.85±0.16  3Hrs 0.87±0.090  4Hrs 1.1±0.072  24Hrs 1.22±0.09  **GFP**  **DCZ**  Baseline  1,23± 0,078  0.5Hr 1,07± 0,083  1Hr 1,28± 0,14  2Hrs 1,22 ±0,12  3Hrs 1,233± 0,078  4Hrs 1,32± 0,097  24Hrs 1,38 ±0,072 | No | Two Way RM ANOVA | p_interaction_<0.0139  F_interaction_=3.555 | Sidak’s  Gi vs GFP  p_row 1 and 3_ = 0.0043 and 0.0054.  DCZ Gi  Baseline vs 1hr; p=0.0346 | 12 ChR2  12 GFP | NN |
| S6C | before stim opto; 1.57±0.13Hz; after stim opto; 0.5±0.06Hz | Yes | Paired t test | p<0.0001 | NN | 16 | NN |
| S7B | **ArchT**  light OFF 12.9±2.6%  light ON  92.5±2.8%  **GFP**  light OFF.  10±6.5%  light ON  10±5.3% | No | Two Way RM ANOVA | p_interaction_<0.0001  F_interaction_=83.24 | Sidak’s  ArchT  Light OFF vs light ON  p < 0.0001.  GFP  Light OFF vs light ON  p=0.99 | 24 ArchT  8 GFP |  |
| S7C | **ArchT**  light OFF 5.32±0.42s  light ON  4.04±0.32s  **GFP**  light OFF.  3.97±0.31s  light ON  3.745±50.28s | No | Two Way RM ANOVA | p_interaction_<0.0001 | Sidak’s  ArchT  Light OFF vs light ON  p < 0.0001.  GFP  Light OFF vs light ON  p=0.398 | 23 ArchT  17 GFP | NN |
| S7D  Males | **ArchT**  light ON  1.4±0.42g  light OFF.  3.2±0.5g | No | Wilcoxon | p=0.0312 | NN | 7 | NN |
| S7D  Females | **ArchT**  light ON  1.38±0.17g  light OFF.  3.5±0.4g | No | Wilcoxon | p=0.0001 | NN | 17 | NN |
| S7E  Males | **ArchT**  light ON  80±4.4%  light OFF. 11.43±4.6% | No | Wilcoxon | p=0.0156 | NN | 7 | NN |
| S7E  Females | **ArchT**  light ON  89.41±3.8%  light OFF. 13.53±3.2% | No | Wilcoxon | p<0.0001 | NN | 17 | NN |
| S7F  Males | **ArchT**  light ON  3.7±0.6s  light OFF  5±0.63s | Yes | Paired t test | p=0.0014 | NN | 7 | NN |
| S7F  Females | **ArchT**  light ON  4.2±0.39s  light OFF  5.5±0.55s | Yes | Paired t test | p<0.0001 | NN | 16 | NN |
| S8B | ArchT  Light OFF  3.9±0.5  Light ON  3.9±0.46 | Yes | Paired t test | p=0.7759 | NN | 8 | 17 |
| S9B | **ChR2**  light ON  16.7±6.15%  light OFF. 93.3±6.7%  **GFP**  light ON  98±2%  light OFF  100±0% | No | Two Way RM ANOVA | p_interaction_<0.0001  F_interaction_=105.3 | Sidak’s  Light OFF vs Light ON ChR2  p<0.0001  Light OFF vs Light ON GFP  p=0.989 | 6 ChR2  10 GFP | NN |
| S9D | ChR2  **5hz/5ms**  light OFF  2.6±0.19g  light ON.  6.8±0g  **5hz/100ms**  light OFF  2±0.19g  light ON  0.58±0.05g | No | Two Way RM ANOVA | p_interaction_<0.0001  F_interaction_=290.1 | Sidak’s  Light OFF vs Light ON ChR2. 5Hz-5ms  p<0.0001  Light OFF vs Light ON ChR2. 5Hz. 100ms  p=0.0007 | 5 | NN |
| S9E | ChR2  **5hz/5ms**  *light ON. 9.83±0.54s*  *light OFF 6.5±0.6s*  **5hz/100ms**  *light ON 2.8±0.11s*  *light OFF. 4±0.1s* | No | Two Way RM ANOVA | p_interaction_<0.0001  F_interaction_=93.96 | Sidak’s  Light OFF vs Light ON ChR2. 5Hz-5ms  p<0.0001  Light OFF vs Light ON ChR2. 5Hz. 100ms  p=0.0142 | 5 | NN |
| S9G | ChR2 :  Light OFF; 5.1±0.63 spikes;  Light ON; 5.35±0.66 spikes. | Yes | Paired t test | p=0.2874 | NN | 10 | 20 |
| S9H | GFP:  Light OFF; 5.4±0.92 spikes;  Light ON; 5.7±1.1 spikes. | Yes | Paired t test | p= 0.358 | NN | 3 | 15 |
| S11B | Naive  1.42±0.46  GFP  2.58±0.79  ChR2  7.42±0.83 | No | Kruskal  Wallis | p<0.0042  Kruskal Wallis Stat =8.03 | Dunn’s  cFOS vs cFOS ChR2  p=0.0179 | 4 | 12 |
| S11C | Naive  0.42±0.32  GFP  0.75±0.25  ChR2  2.92±0.48 | No | Kruskal  Wallis | p=0.0088  F =7.8 | Dunn’s  cFOS Pax2 vs cFOS Pax 2 ChR2  p=0.0287 | 4 | 12 |
| S11D2 | 5-HT2a/glut  0.2±0.02  5-HT2a/gaba. 0.73±0.013 | Yes | Paired t test | p<0.0001 | NN | 2 | 6 |
| S11E2 | 5-HT2a/Pax2  0.54+/-0.04  5-HT2a/Tlx3  0.013+/-0.004 | Yes | Paired t test | p<0.0001 | NN | 3 | 9 |
