## Supplementary material for "Nucleus raphe magnus serotonin neurons bidirectionally control spinal nociceptive transmission in mice": Table S3

| Figure N° | Mean  ±SEM | Normality test  Shapiro-wilk test | Statistic method | Main effect | Post hoc | Number of mice | Number of samples |
| --- | --- | --- | --- | --- | --- | --- | --- |
| 1B  DCZ | **DCZ**  baseline 1.367±0.02305  0.5Hr 1.017±0.06907  1Hr  0.8±0.06138  2Hr  0.9±0.06019  3Hr 1.217±0.07114  4Hr  1.2±0.06370  24Hr 1.517±0.06246  **Saline**  baseline  1.3±0.01  0.5Hr  1.22±0.118  1Hr  1.289±0.1087  2Hr  1.42±0.08  3Hr  1.478±0.061  4Hr  1.34±0.056  24Hr  1.36±0.030 | No | Two Way ANOVA | p_interaction_=0.0089  F_interaction_=6.378 | Sidak’s  Gq DCZ  Baseline vs 30min p=0.0051  Baseline vs 1hr  p<0.0001  Baseline vs 2hrs  p<0.0001  DCZ vs Saline  30min p=0.044  1hr  p<0.0001  2hrs  p=0.0002  3hrs  p=0.0107 | 24 Gq DCZ  18 Gq Saline | NN |
| 1D  ChR2 | **ChR2**  Light OFF  2.045±0.2076  Light ON  0.7826±0.09960  **GFP**  Light OFF  1.959±0.1854  Light ON  1.867±0.1869 | No | Two Way RM ANOVA | p_interaction_<0.0001  F_interaction_=48.98 | Sidak’s  Baseline vs opto (ChR2)  p<0.0001  Baseline vs opto (GFP)  p=0.87 | ChR2 17  GFP 24 | NN |
| 1H | **ChR2**  Light OFF  10.84±2.147  Light ON  14.00±2.164  **GFP**  Light OFF  8.149±1.131  Light ON  8.252±1.132 | No | Two Way RM ANOVA | p_interaction_=0.0006  F_interaction_=13.13 | Sidak’s  Baseline vs opto (ChR2)  p<0.0001  Baseline vs opto (GFP)  p=0.87 | 5 (Chr2)  9 (GFP) | 35 (ChR2)  30 (GFP) |
| 1I | **Light OFF**  Stim 1: 7.5±1.91  Stim 2 7.67±1.61  Stim 3 7.83±1.851  Stim 4 10.17±1.904  Stim 5 10.3±2.40  Stim 6 12.67±2.47  Stim 7 14.67±2.641  Stim 8 14.83±2.414  Stim 9 15.33±2.728  Stim 10 18.17±2.5  **Light ON**  Stim 1: 8.83±2.761  Stim 2 10.17±3.32  Stim 3 11±2.5  Stim 4 12.67±3.26  Stim 5 12±3.28  Stim 6 13.67±3.01  Stim 7 12.83±3.05  Stim 8 14.33±3.658  Stim 9 14.83±4.1  Stim 10 15.83±3.34 | No | Two Way RM ANOVA | p_interaction_=0.9983  F_interaction_=0.38 | Sidak’s  Light on vs Light OFF, p>0.05 | NN | 6 |
| 1J | **Light OFF** 44.17±6.54  **Light ON**  37.83±18.01 | Yes | Paired t-test | p=0.69 | NN | NN | 6 |
| 2B | 5-HT3/GABA  0.097±0.013  5-HT3/Glut  0.64±0.05 | Yes | Unpaired t-test. | p<0.0001 | NN | 3 | 9 |
| 2D | **Vehicle**  light OFF  1.9±0.25g  light ON : 0.86±0.11g  **Granisetron**  light OFF  1.6±0.21g  Light ON : 2.23±0.45g | No | Two Way RM ANOVA | p_interaction_<0.0001  F_interaction_=23.12 | Sidak’s  Saline  Baseline vs opto (ChR2)  p=0.002  Granisetron  Baseline vs opto (ChR2)  p=0.04 | 12 | NN |
| 3B | **Gi**  **DCZ**  Baseline 1.35±0.078  0.5Hr 1.15±0.176  1Hr 0.7167±0.1114  2Hrs 0.85±0.16  3Hrs 0.87±0.090  4Hrs 1.1±0.072  24Hrs 1.22±0.09  **Saline**  Baseline 1.583±0.1140  0.5Hr 1.683±0.1192  1Hr 1.567±0.1453  2Hrs 1.550±0.1234  3Hrs 1.650±0.1306  4Hrs 1.633±0.1176  24Hrs 1.533±0.1082 | No | Two Way RM ANOVA | p_interaction_<0.0077  F_interaction_=3.901 | Sidak’s  Gi DCZ  Baseline vs 1hr p=0.034  DCZ vs Saline  30min p=0.0212  1hr  p=0.0001  2hrs  p=0.0028  3hrs  p<0.0001  4hrs  p=0.0011 | 12 | NN |
| 3D | **ArchT**  Light OFF  2.32±0.17g  Light ON  0.98±0.09g  **GFP**  Light OFF  1.58±0.18g  Light ON  1.42±0.16g | No | Two Way RM ANOVA | p_interaction_<0.0001  F_interaction_=29.53 | Sidak’s  Baseline vs opto (ArchT)  p<0.0001  Baseline vs opto (GFP)  p=0.85 | ArchT 27  GFP 14 | NN |
| 3H | **ArchT**  Light OFF  4.8±1.38  Light ON  10.2±2.31  **GFP**  Light OFF  7.5±2.9g  Light ON  8±2.9g | No | Two Way RM ANOVA | p_interaction_=0.0107  F_interaction_=7.77 | Sidak’s  Baseline vs opto (ArchT)  p=0.0001  Baseline vs opto (GFP)  p=0.996 | 10 ArchT  3 GFP | 16  7 |
| 3I | **Light OFF**  Stim 1 1.3±0.71  Stim 2 1.14±0.55  Stim 3 1.43±0.48  Stim 4 2±0.65  Stim 5 2.14±1.01  Stim 6 2.86±1.06  Stim 7 2.86±1.26  Stim 8 2.86±0.89  Stim 9 3.14±1.056  Stim 10 2.86±0.829    **Light ON**  Stim 1 4±1.04  Stim 2 5.1±1.14  Stim 3 6.29±1.15  Stim 4 6.43±1.27  Stim 5 7.29±0.89  Stim 6 8.43±1.21  Stim 7 8±1.1  Stim 8 8±1.53  Stim 9 8.14±1.06  Stim 10 9±1.41 | Yes | Two-Way RM ANOVA | P_rank of stim_=0.0024  P_opto_=0.0033  F_rank of stim_ =10.12  F_opto_=0.0033 | Sidak’s  p_row2.4.7.8._=0.0123 ; 0.0128 ; 0.01 ; 0.0161  p_row3.5.6.9.10_=0.0045 ; 0.0025 ; 0.0048 ; 0.0058.  p_row10_= 0.0040 | NN | 7 |
| 3J | Light OFF  9.71±1.71  Light ON  30.71±4.61 | No | Wilcoxon  Test | p=0.0156 | NN | 3 | 7 |
| 4H | 5-HT2c/Glut  0.54±0.03  5-HT2c/gaba  0.25±0.02 | Yes | Paired t test | p=0.0002 | NN | 3 | 7 |
| 4J | Tomato/tlx3  0.21±0.03  Tomato/Pax2  0.14±0.02 | Yes | Paired t-test | p=0.03 | NN | 3 | 12 |
| 4L | Control  1.25±0.11g  DMSO  1.27±0.14g  NaCl  1.5±0.14g  Methysergide 0.63±0.25g. RS102221 : 0.27±0.08g  Ketanserin  1.33±0.17g | No | Kruskal-Wallis | p<0.0001 | Dunn’s  p_control vs RS10221_=0.0009 | NaCl =6  DMSO =11  Control=17  Methysergid=6  RS10221=11  Ketanserin =6 | NN |
| 5B | ChR2  Light OFF  1.93±0.16g Light ON  4.08±0.42g  GFP  Light OFF  1.96±0.19g Light ON  1.94±0.21g | No | Two-Way RM ANOVA | P_interaction_ <0.0001  F_interaction_ =33.96 | Sidak’s  Baseline vs opto (ChR2)  p=0.0001  Baseline vs opto (GFP)  p=0.9998 | ChR2 24  GFP 14 | NN |
| 5D | ChR2.  light OFF 5.5±0.7  light ON 4.2±0.6  GFP  light OFF 8±1.6  light ON 8.4±1.7 | No | Two-Way RM ANOVA | P_interaction_ <0.0001  F_interaction_ =27.68 | Sidak’s  Baseline vs opto (ChR2)  p=0.0001  Baseline vs opto (GFP)  p=0.45 | ChR2 14  GFP 3 | 31  15 |
| 5E | **Light OFF**  Stim 1 6.33±1.48  Stim 2 7.92±1.75  Stim 3 8.83±2.34  Stim 4 11.75±3.092  Stim 5 12.76±3.25  Stim 6 15.33±3.9  Stim 7 17.3 ±3.79  Stim 8 17.25±3.8  Stim 9 18.7±4.19  Stim 10 18.75±3.36  **Light ON**  Stim 1 8.33±2.05  Stim 2 9±2.17  Stim 3 8.08±1.73  Stim 4 10.17±2.28  Stim 5 9.5±2.44  Stim 6 10.5 ±1.87  Stim 7 12.92±2.46  Stim 8 11.5±2.162  Stim 9 12.5 ±2.34  Stim 10 13.08±2.18 | No | Two Way RM ANOVA | p_opto_=0.69  F=0.16  p_rank of stim_<0.0001  F=20.92 | Sidak’s  p_light off vs light on_ >0.05  p _light ON_ _row10._=0.0049 | 3 | 12 |
| 5F | **ChR2**  Light OFF  71.50±17.91  Light ON  23.00±6.592  **GFP**  Light OFF  28.85±9.134  Light ON  23.38±5.563 | No | Two Way RM ANOVA | P_interaction_ <0.0255  F_interaction_ =5.7 | Sidak’s  Baseline vs opto (ChR2)  p=0.0044  Baseline vs opto (GFP)  p=0.9875 | 3 | 12 |
| 6A | **Vehicle**  Light OFF  1.42±0.14g  Light ON  2.13±0.16g  **Methysergide**  Light OFF  0.63±0.25g Light ON  0.79±0.4g | No | Two Way RM ANOVA | p_interaction_=0.0089  F_interaction_=10.47 | Sidak’s  p_vehicle vs opto_=0.0006 | 6 | NN |
| 6B | **Vehicle**  Light OFF  1.27±0.14g  Light ON  1.93±0.23g  **RS10221**  Light OFF  0.27±0.08g  Light ON  0.23±0.07g | No | Two Way RM ANOVA | p_interaction_=0.039  F_interaction_=5.05 | Sidak’s  p_vehicle vs opto_=0.038 | 9 | NN |
| 6D | **Vehicle**  Light OFF  1.4±0.13g  Light ON  2.29±0.21g  **Ketanserin**  Light OFF  1.33±0.17g  Light ON  1.56±0.26g | No | Two Way RM ANOVA | p_interaction_=0.0007  F_interaction_=22.95 | Sidak’s  p_vehicle vs opto_=0.0001 | 6 | NN |
| 6F | **Vehicle**  Light OFF  1.67±0.14g  Light ON  2.95±0.33g  **Granisetron**  Light OFF  1.17±0.21g  Light ON  2.93±0.43g | No | Two Way RM ANOVA | p_interaction_=0.4032  F_interaction_=0.7243 | Sidak’s  p_vehicle vs opto_=0.0003  p_granisetron vs opto_<0.0001 | 13 | NN |
