## Supplementary material for "Nucleus raphe magnus serotonin neurons bidirectionally control spinal nociceptive transmission in mice": Table S2

Table S2 : List of 5-HTRs antagonists.

| **Antagonist** | **Reference** | **Final doses in 10µL** | **Solvent** | **Target** |
| --- | --- | --- | --- | --- |
| Methysergide | Tocris (1064) | 20 µg in 10µL | NaCl 0.9% | 5-HT_1/2_ |
| Ketanserin | Tocris (0908) | 20 µg in 10µL | NaCl 0.9% | 5-HT2_A_ |
| RS1002221 | Tocris(1050) | 10 µg in 10µL | DMSO 1% | 5-HT2_C_ |
| Granisetron | Tocris (2903) | 15 µg in 10µL | DMSO 1% | 5-HT3 |
