## Supplementary material for "Nucleus raphe magnus serotonin neurons bidirectionally control spinal nociceptive transmission in mice": Table S1

Table S1: Viruses used for optogenetic and chemogenetic.

| **Protein** | **Virus** | **Ref** |
| --- | --- | --- |
| Archaerhodospin (ArchT) | AAV9-CAG-flex-ArchT-GFP | UNC Vector Core - AV6222B |
| Channelrhodospin (ChR2) | AAV-EF1a-DIOhChR2(H134R)-EYFP-WPRE- | Vector core - AV4313Y or Addgene - 37082 |
| GFP (control) | AAV9.CAG-Flex-eGFP-WPRE | Addgene - 51502 |
| DREADDs - hM4(Gi) | AAV-hsyn-DIO-hM4D(Gi)-mCitrine | Addgene – 50455 |
| DREADDS - hM3(Gq) | -hSyn-DIO-hM3D(Gq)-mCitrine | Addgene – 50454 |
