## Supplementary figures and images for "Nucleus raphe magnus serotonin neurons bidirectionally control spinal nociceptive transmission in mice"

### Fig Sup 9

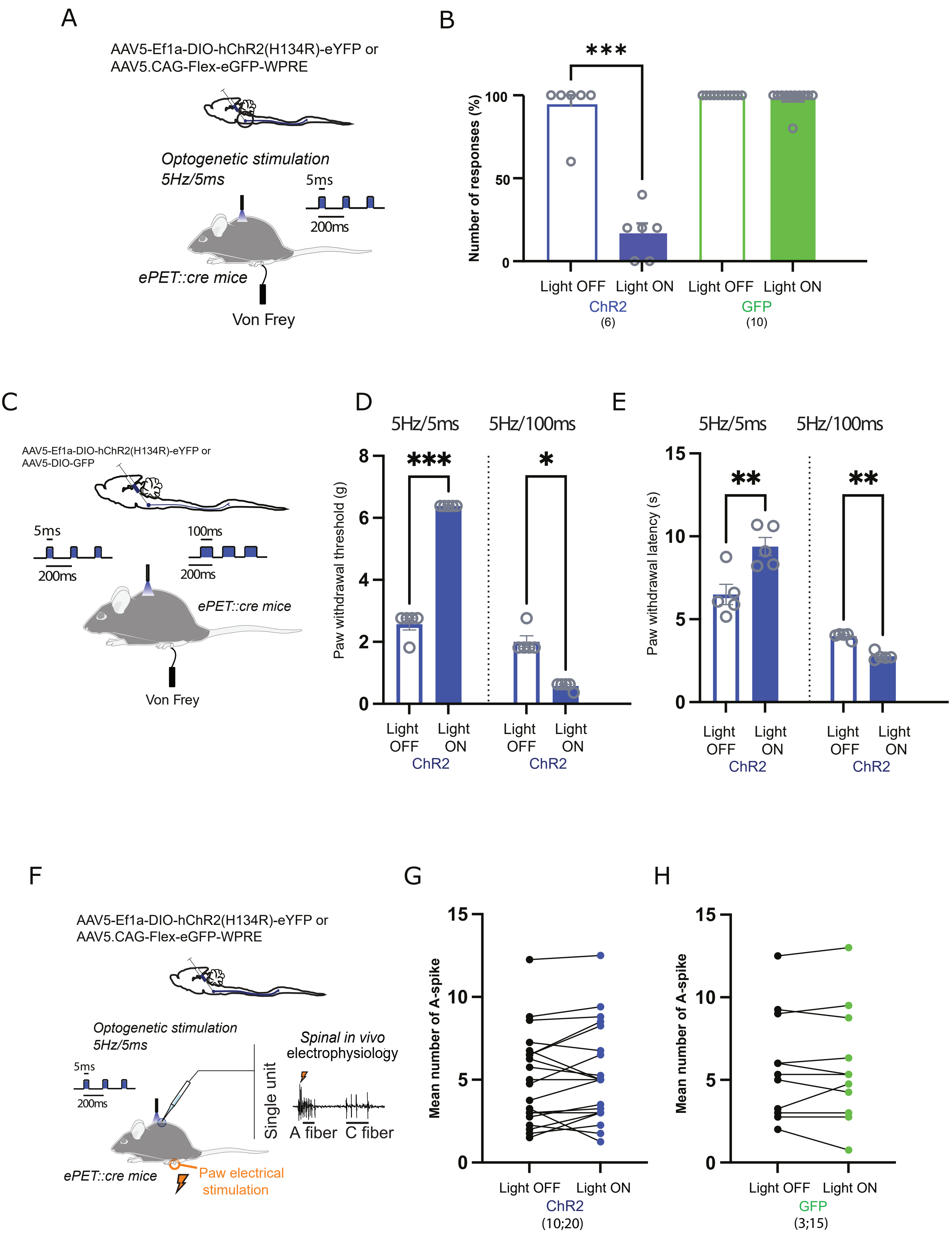

### Sup Fig 1

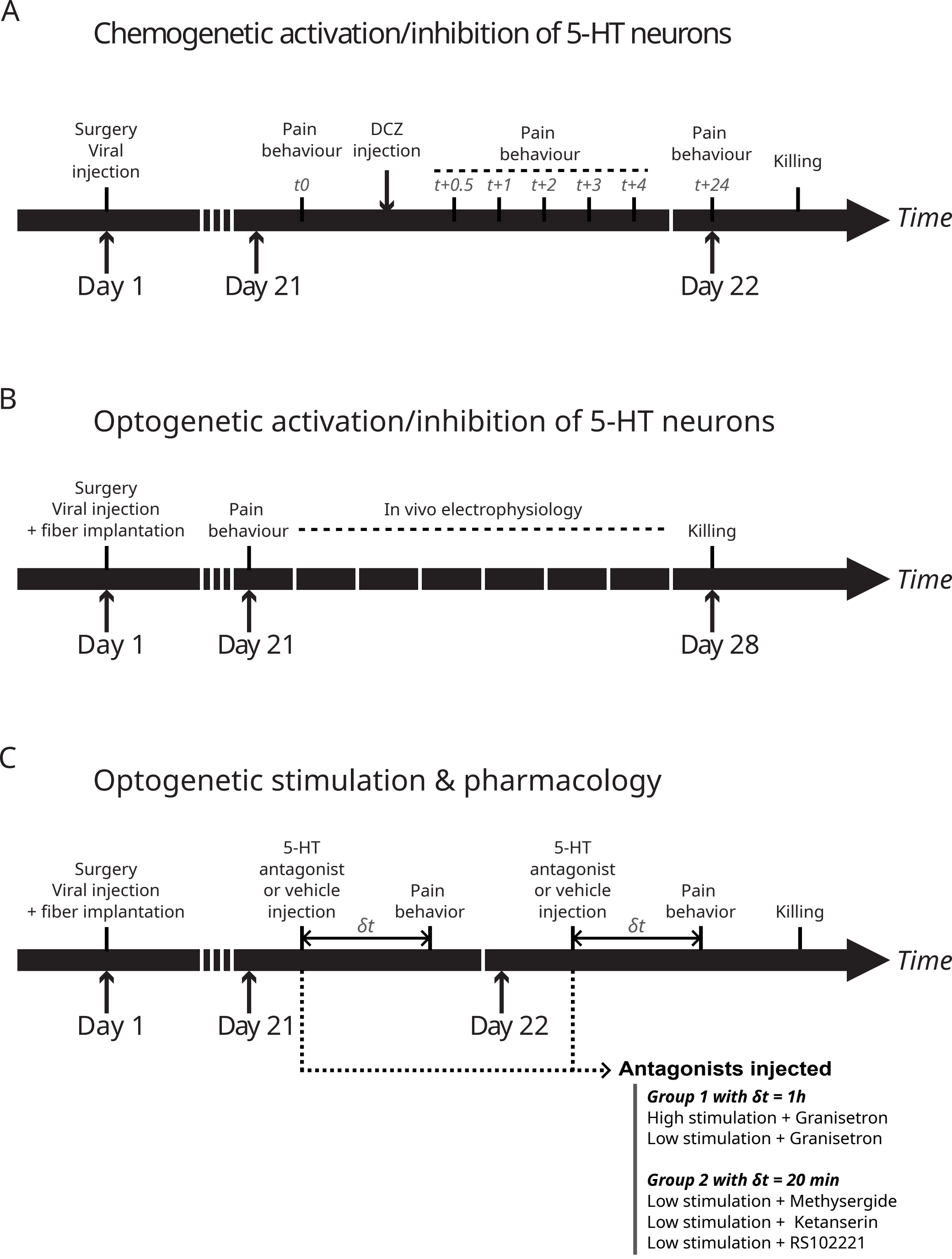

### Sup Fig 2

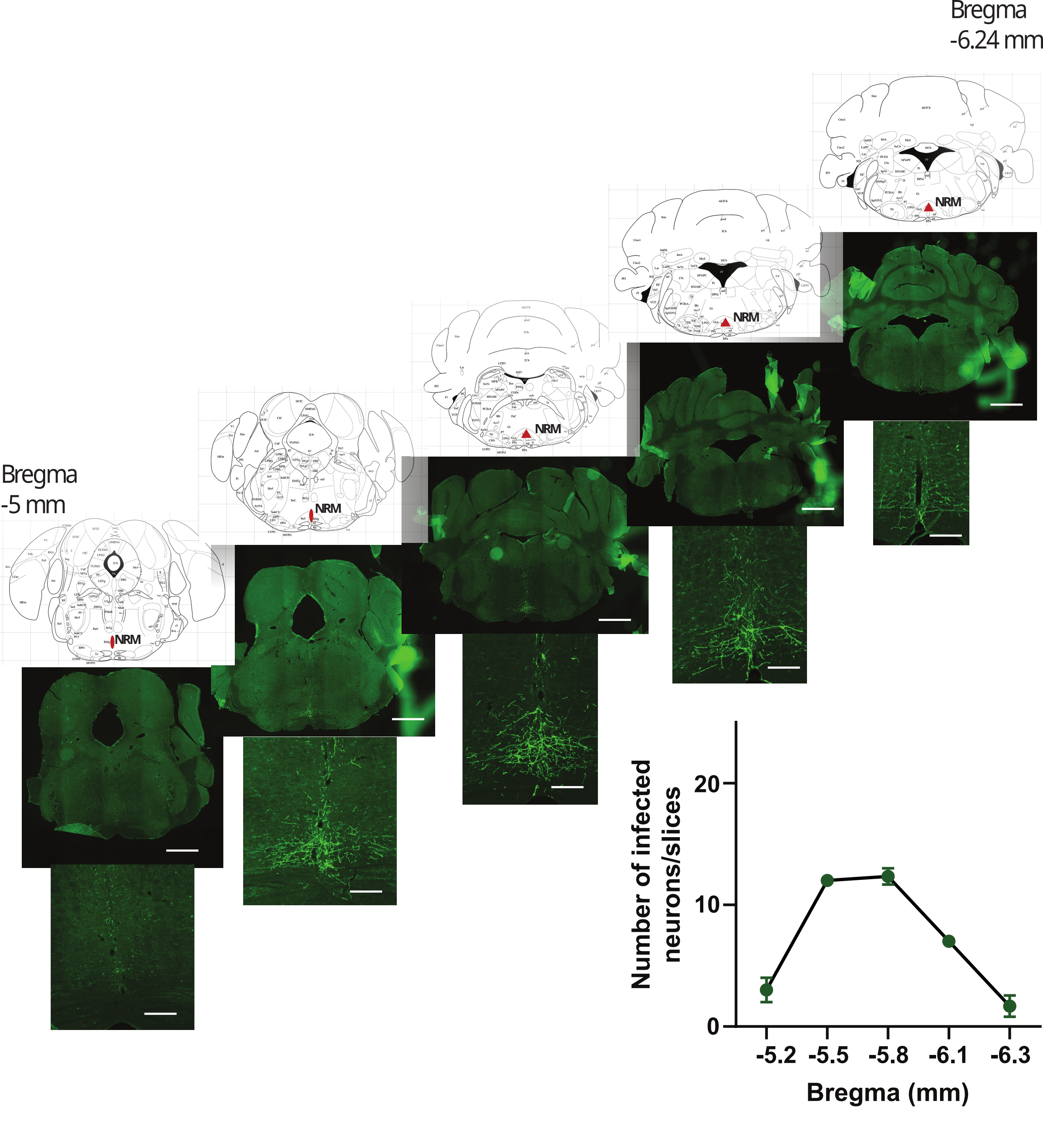

### Sup Fig 3

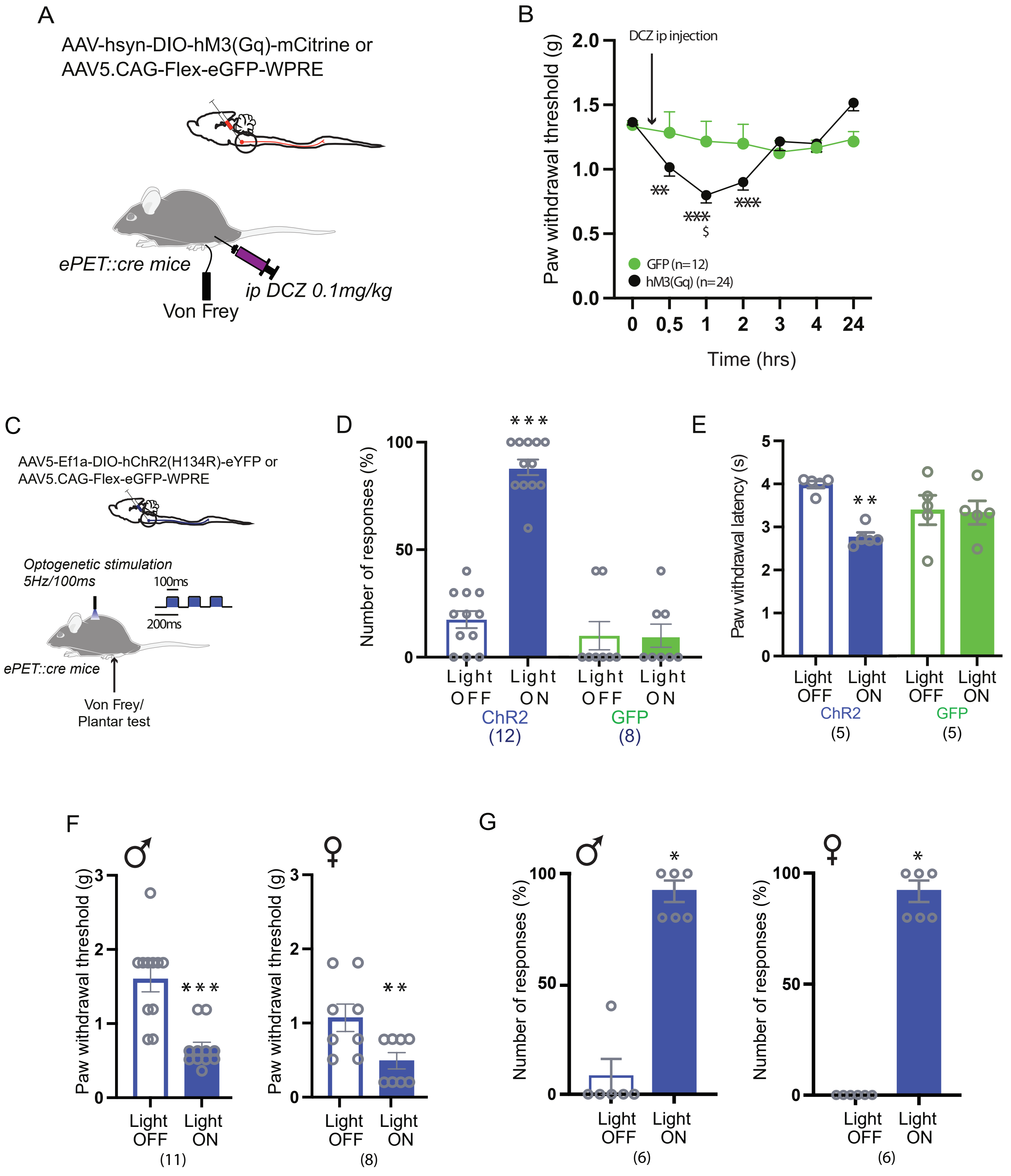

### Sup Fig 4

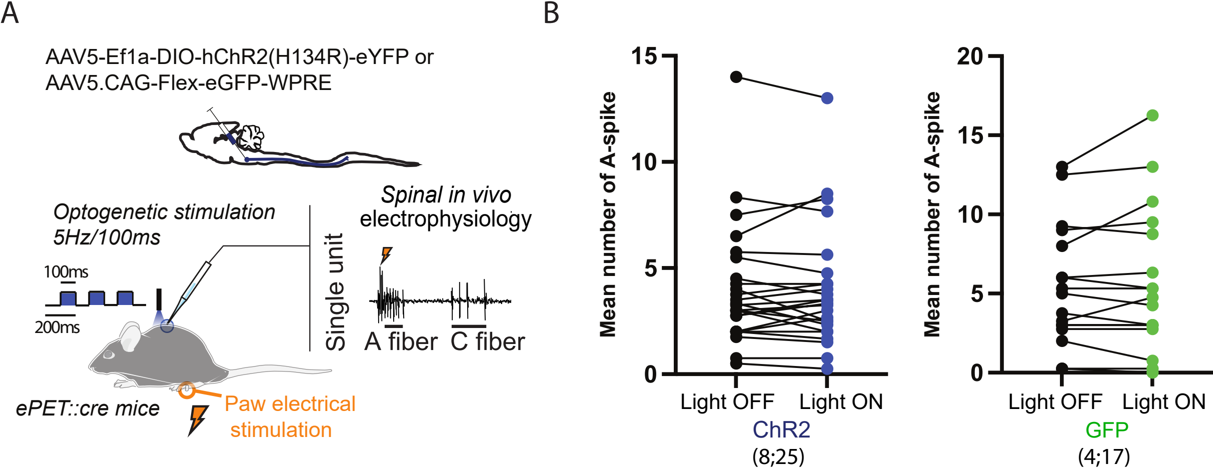

### Sup Fig 5

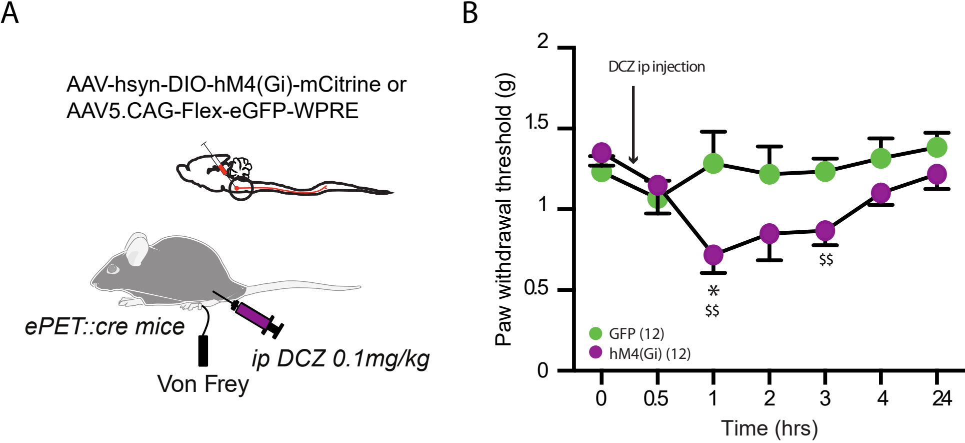

### Sup Fig 6

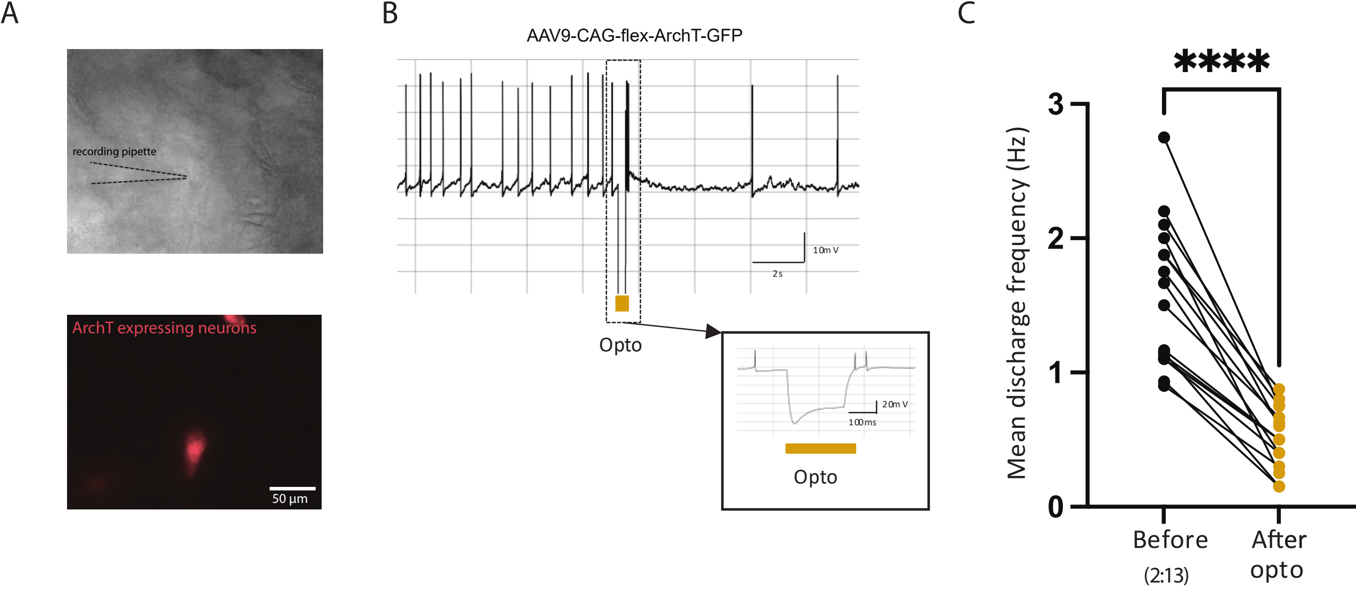

### Sup Fig 7

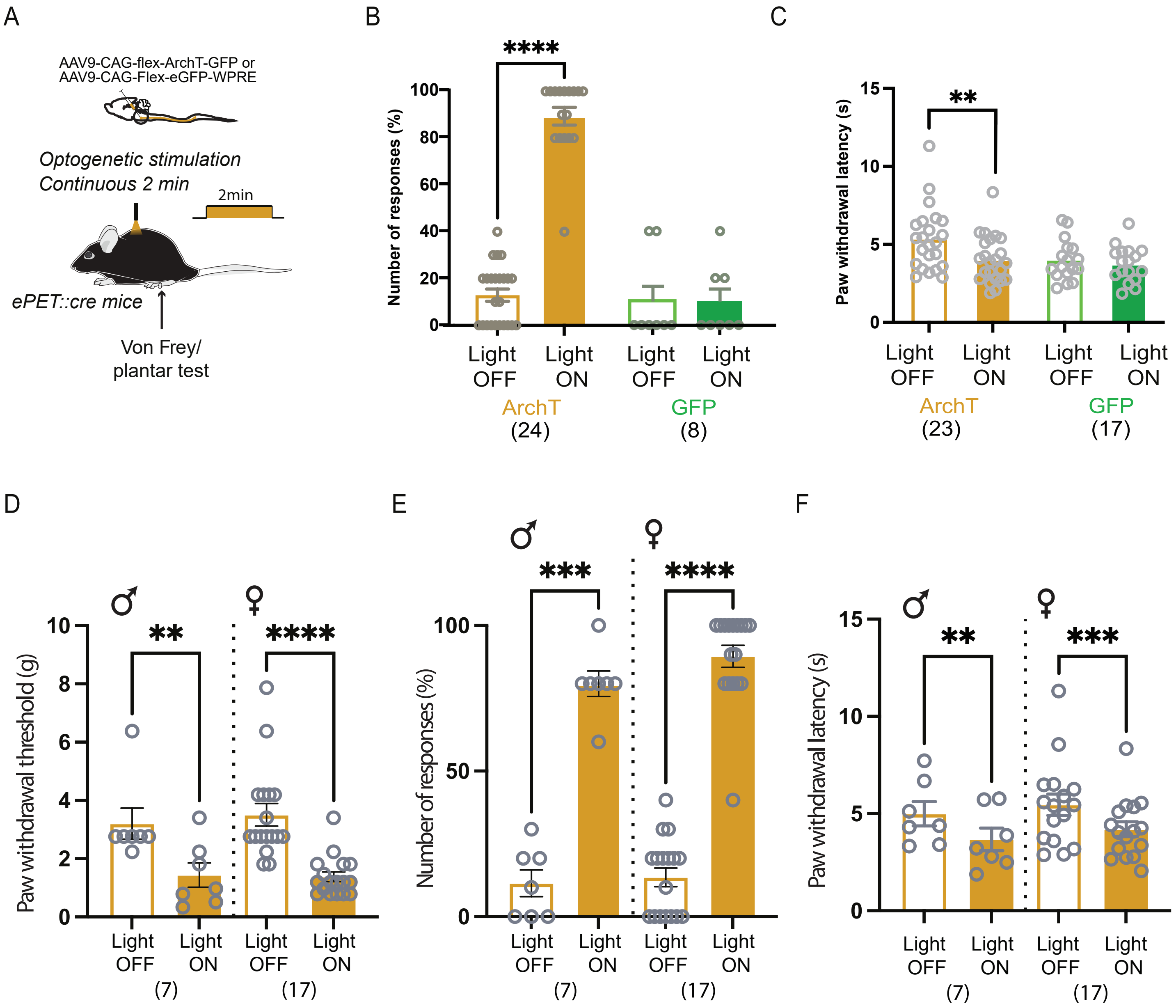

### Sup Fig 8

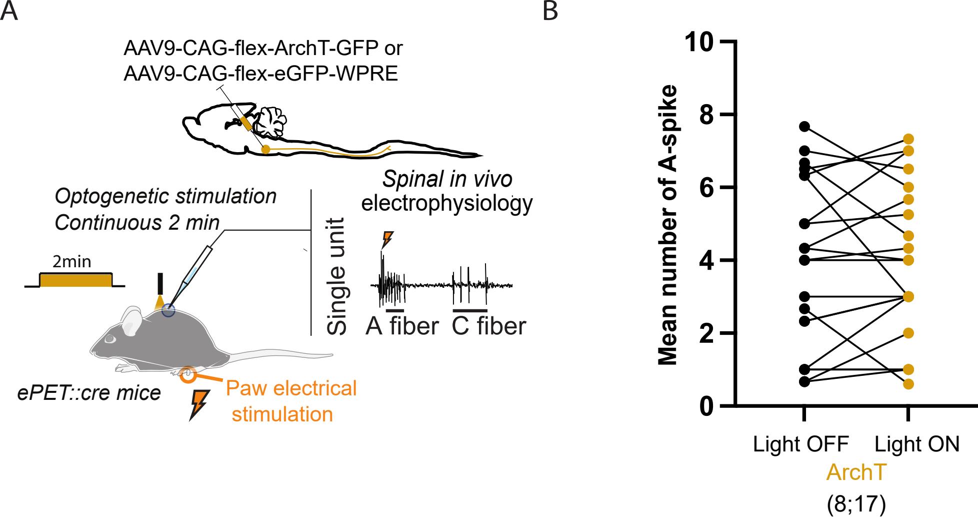

### Sup Fig 10

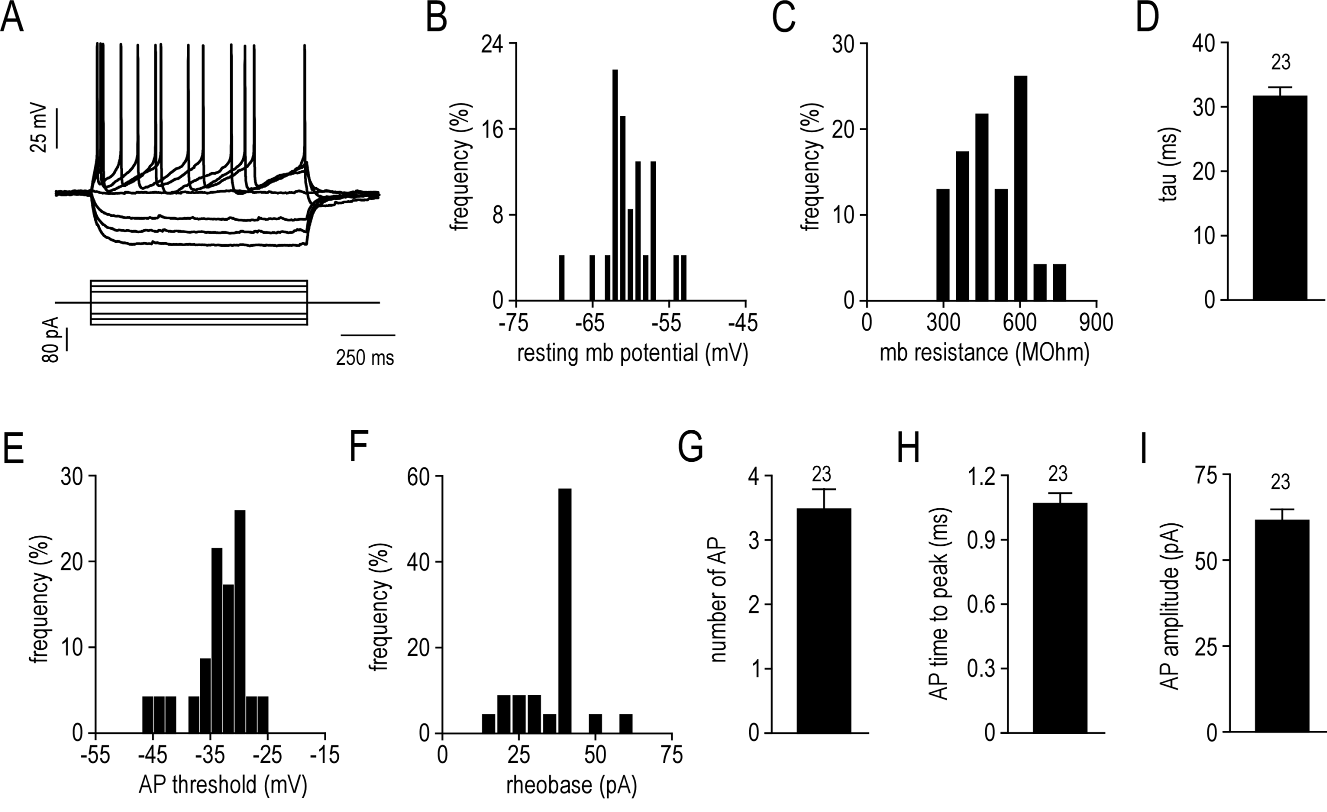

### Sup Fig 11

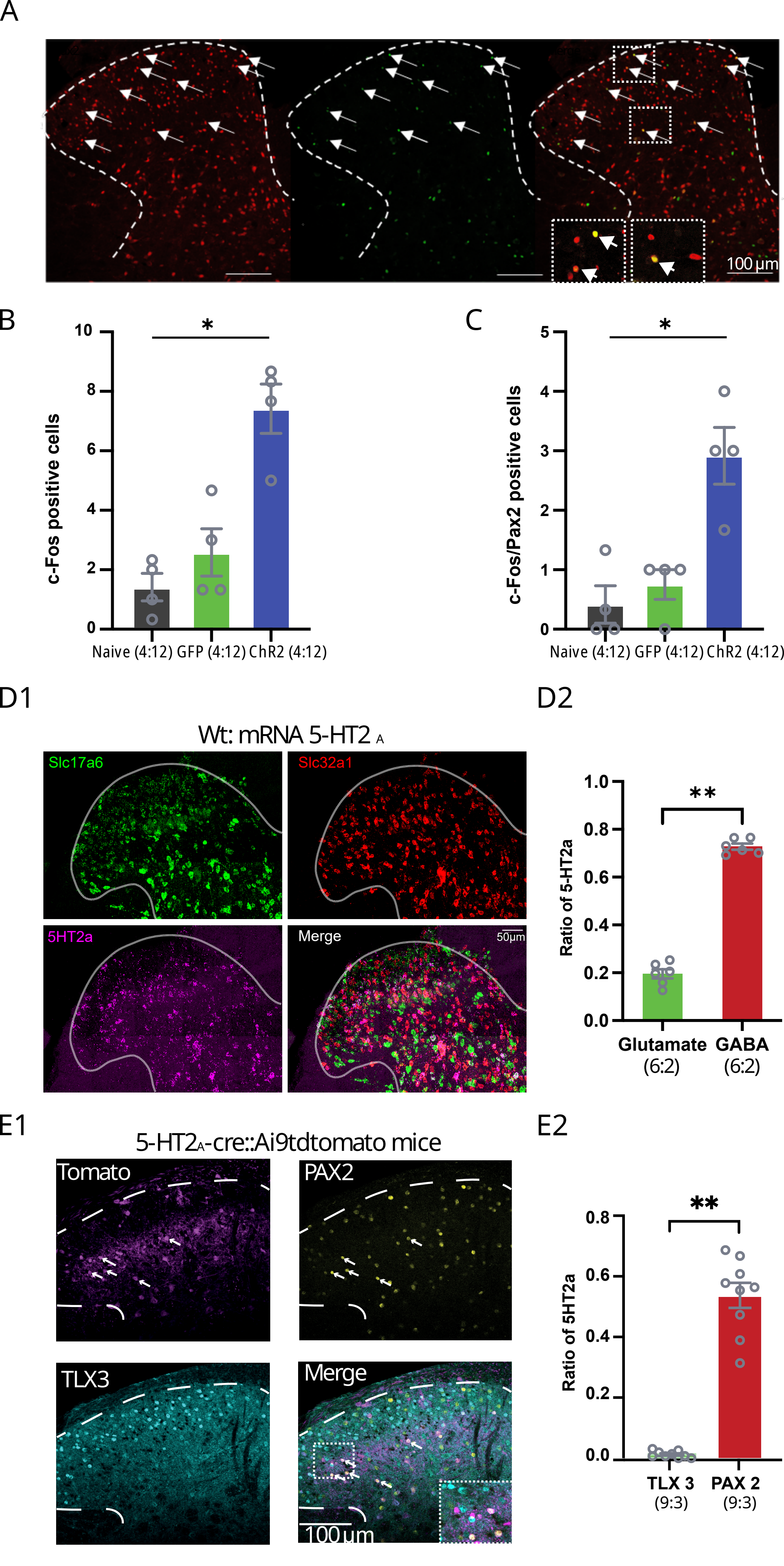

### Sup Fig 12

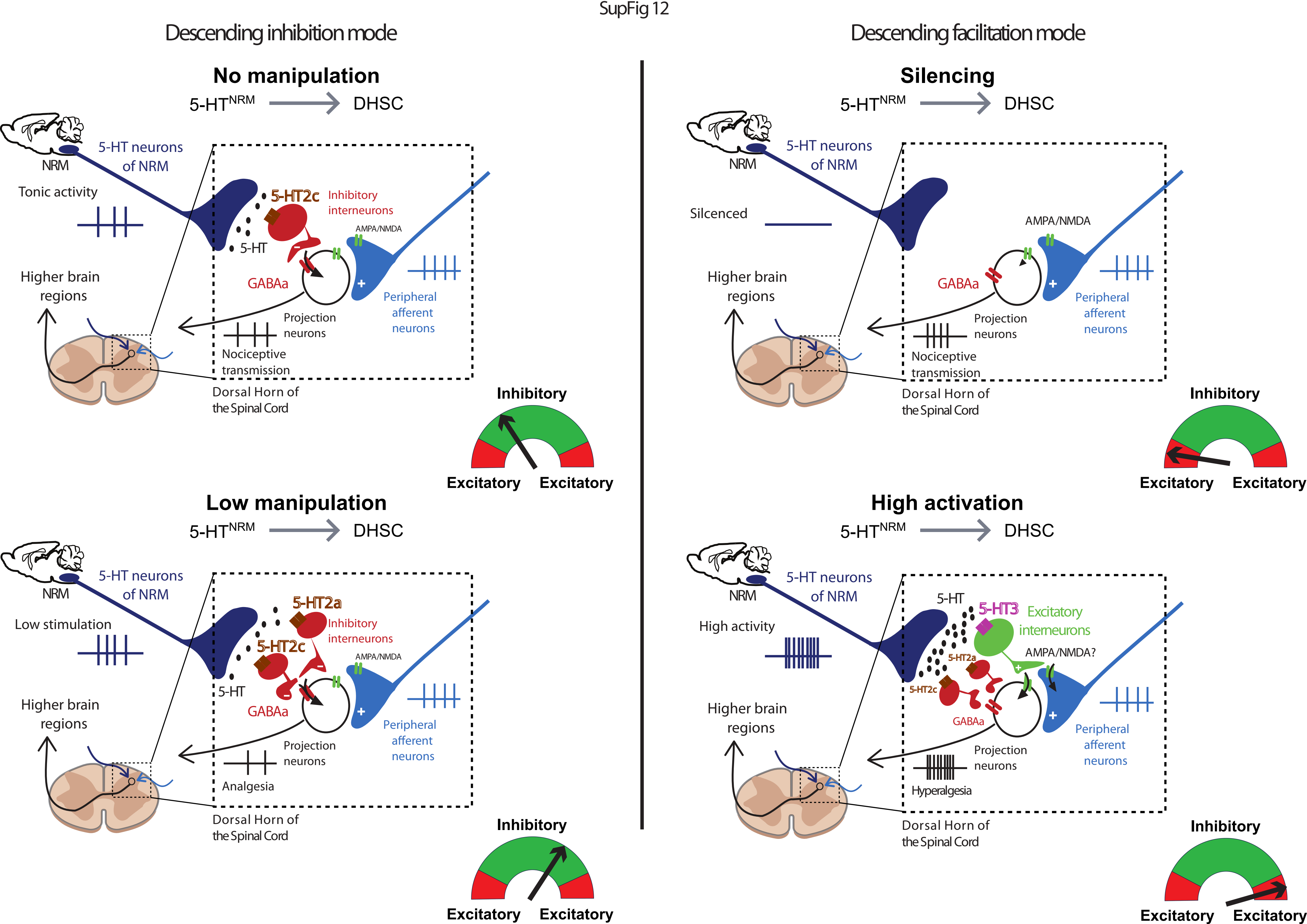
